# Slow-wave sleep alters the stability landscape of synaptic-weight space allowing life-long learning

**DOI:** 10.1101/2025.11.10.687549

**Authors:** Oscar C. González, Ryan Golden, Erik Delanois, Bruce L. McNaughton, Maxim Bazhenov

**Affiliations:** Department of Integrative Physiology, University of Colorado Boulder; School of Medicine, University of California, San Diego; Neurosciences Graduate Program, University of California, San Diego; Department of Computer Science and Engineering, University of California, San Diego; Department of Neurobiology and Behavior, University of California, Irvine; Canadian Centre for Behavioral Neuroscience, University of Lethbridge

## Abstract

Sleep replay - the reactivation of memory traces during slow-wave sleep - is widely held to stabilize memories and reduce interference, yet exactly how replay reorganizes synaptic-weight space to preserve existing memories while incorporating new ones remains unclear. Here, we use a biophysically realistic network model to probe the synaptic-weight dynamics underlying this process. We find that replay drives synaptic weights toward stable configurations - synaptic attractors - that jointly support both old and new memories. Hippocampus-driven interactions between sharp-wave ripples and cortical slow waves guide this reorganization, allowing recently acquired memories to be incorporated without degrading prior ones. These results reveal a mechanistic and geometric framework for memory consolidation: sleep does not passively protect memories, but actively sculpts synaptic-weight space into attractor configurations from which forgetting requires escaping a stability basin.

**SIGNIFICANCE STATEMENT:** Storing, processing, and retrieving information underpins intelligent behavior. Sleep extracts invariant features from prior experience, promoting the emergence of explicit knowledge and insight. Yet despite abundant empirical findings, our understanding of how sleep reshapes memory representations at the level of synaptic organization remains limited. Here we present a novel framework that describes how memories are encoded in synaptic-weight space and how sleep dynamics reorganize synaptic landscape. These results advance our understanding of how the brain solves core problems of lifelong learning.

## INTRODUCTION

Continual learning is a foundation of human intelligence. To survive in constantly changing environments, brains must continuously encode new memories without erasing existing ones - minimizing interference while allowing prior knowledge to scaffold new learning (Brod et al., 2016). Our limited understanding of how the brain accomplishes this is underscored by ongoing attempts to achieve scalable continual learning in artificial neural networks (ANNs), which suffer severe retroactive interference known as catastrophic forgetting (Mccloskey and Cohen, 1989; McClelland et al., 1995; French, 1999; Hayes et al., 2021; Kudithipudi et al., 2022).

Sleep has long been hypothesized to play a central role in memory consolidation (Walker and Stickgold, 2004; Ji and Wilson, 2007; Lewis and Durrant, 2011). Two mechanisms are thought to be critical: spontaneous replay of memory traces and the unsupervised synaptic plasticity it drives (Wilson and McNaughton, 1994; Stickgold and Walker, 2007; Wei et al., 2016). Using biophysical models of the thalamocortical network, we previously showed that coordinated replay of recently learned and older memories during slow-wave sleep enables networks to form orthogonal memory representations, allowing overlapping neuronal populations to store competing memories without interference (Wei et al., 2016; Wei et al., 2018; Gonzalez et al., 2020; Wei et al., 2020; Golden et al., 2025). Related work demonstrated that sleep-like replay in ANNs similarly mitigates catastrophic forgetting and improves generalization (Tadros et al., 2020b; Tadros et al., 2020a; Tadros and Bazhenov, 2022; Tadros et al., 2022; Delanois et al., 2023; Bazhenov et al., 2024b, a).

Yet a fundamental question remains: how does sleep-driven plasticity reorganize synaptic-weight space? What configurations does it move the system toward, and why do those configurations support multiple memories without interference? Both awake learning and sleep consolidation modify synaptic weights, but the extreme dimensionality of this space has hindered progress in identifying the principles that govern these dynamics. In this study, we use a biophysical model of the canonical thalamocortical circuit to address this challenge directly. We conceptualize learning and sleep as trajectories through an abstract synaptic-weight space, where each point corresponds to a circuit configuration that supports none, one, or multiple memories. Wake learning moves the system through this space under non-autonomous, input-driven dynamics; sleep induces autonomous synaptic dynamics shaped by internal replay. Only a subset of synaptic configurations constitute stable fixed points of the sleep dynamics - synaptic attractors - and we show that coordinated hippocampal sharp-wave ripples and cortical slow oscillations guide the system toward attractors that jointly encode both old and new memories. This provides a mechanistic and geometric account of how the dual hippocampo–cortical memory system, as proposed by systems-consolidation theory (Wilson and McNaughton, 1994; Rasch and Born, 2013), enables continual learning without catastrophic forgetting.

## RESULTS

The paper is organized as follows. First, we revisit some of our previously published results on the role of hippocampus independent replay in preventing catastrophic forgetting after sequential training (Gonzalez et al., 2020), framed from the perspective of network dynamics in synaptic-weight space. Next, we expand those results to the case of hippocampal indexing and show how sleep indexing can set up learning trajectories so that the neocortical network progressively acquires new memories without erasing the old ones. Finally, we generalize these ideas by proposing that continual learning depends on sleep forming new stable memory attractors. These attractors emerge at the intersections of memory manifolds, defined as subspaces of synaptic weights that support different memories.

### Network model design

The network model utilized throughout the study was built upon thalamocortical models developed in our earlier work (Krishnan et al., 2016; Gonzalez et al., 2020). In short (see Methods for details), the basic circuit (Figure 1A) consists of a single cortical layer with excitatory pyramidal cells (PYs) and inhibitory interneurons (INs), and a single thalamic layer with excitatory thalamocortical cells (TCs) and inhibitory reticular interneurons (REs). All neurons were modeled according to the Hodgkin-Huxley formalism, and synaptic connections between cells were set deterministically within a local radius and held at a constant weight value, except for PY-PY synapses. These intracortical synapses were set probabilistically within a local radius with the initial weight values Gaussian distributed (Figure 1B). During awake and sleep states the strength of these synapses was allowed to vary according to the local spike-timing-dependent plasticity (STDP) rules. The model implemented effects of neuromodulators (Krishnan et al., 2016) to enable transitions between awake state, characterized by asynchronous low-frequency firing, and slow-wave sleep (SWS), characterized by slow oscillations (<1 Hz, (Steriade et al., 1993)). In this model, cortical network is responsible for generating the sleep slow oscillations (Timofeev et al., 2000; Bazhenov et al., 2002) however, thalamus contributes to synchronization properties of sleep slow-waves (Lemieux et al., 2014).

**Fig. 1.**
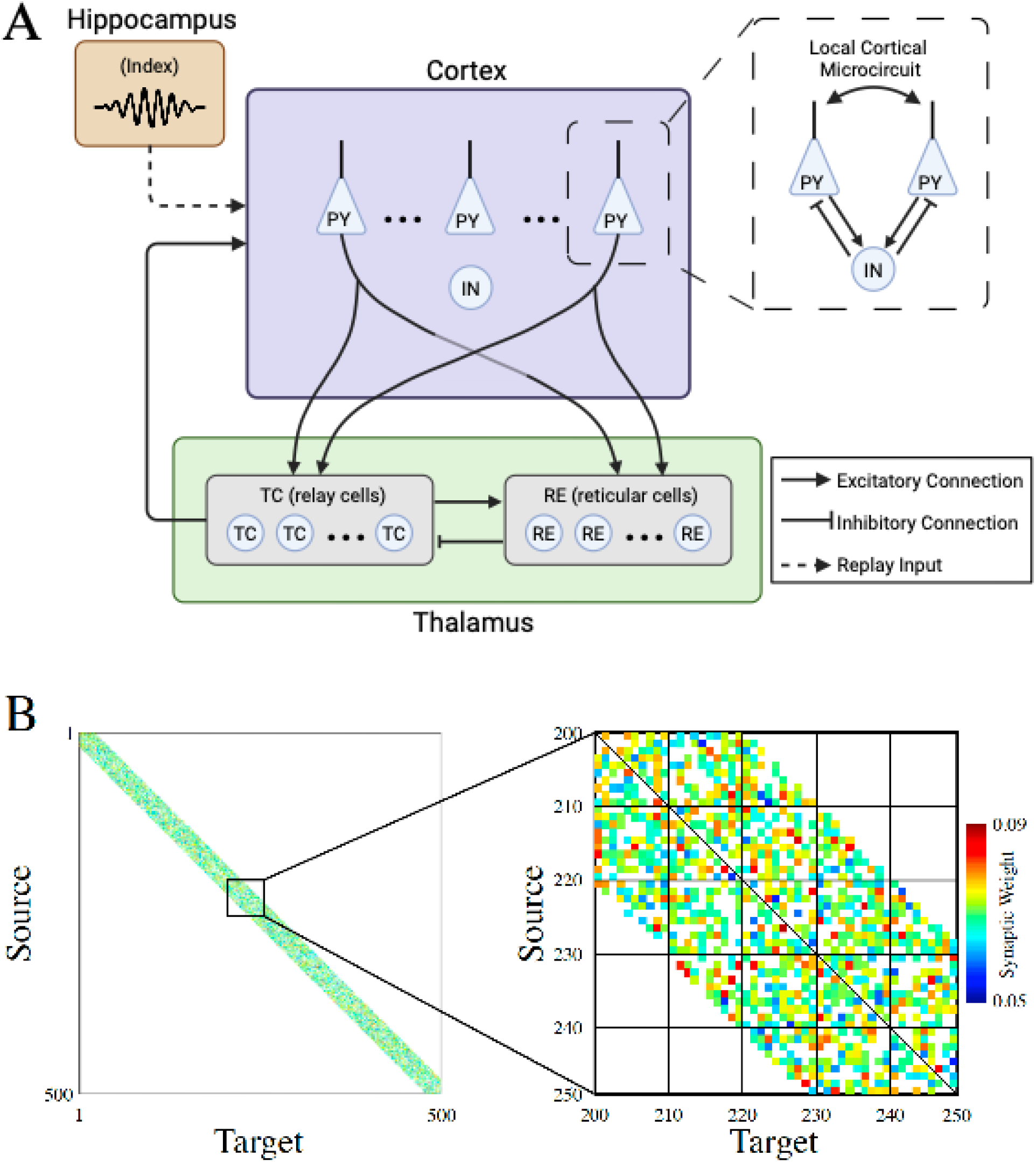
Network Architecture. (**A**) Basic network architecture (PY: excitatory pyramidal neurons; IN: inhibitory interneurons; TC: excitatory thalamocortical neurons; RE: inhibitory thalamic reticular neurons). Excitatory synapses are represented by lines terminating in arrows, while inhibitory synapses are represented by lines terminating in bars. (**B**) The left panel shows the initial synaptic weight matrix for the PY neurons. The color in this plot represents the strength of AMPA connections between PY neurons, with white indicating the absence of a synaptic connection. The right panel shows a zoomed-in view of the top panel for the subregion where training occurs.

### Differential replay of sequence memories during sleep improves recall performance

An example of an experimental simulation paradigm consisting training and testing in awake state and a period of SWS is shown in Figure 2A. First, during the awake state, the network was trained with a single memory - sequence S1 or S2, represented by five sequentially ordered cell groups of ten neurons per group (S1=EDCBA or S2=ABCDA, respectively; see Figure 2B, left and middle). These sequences were chosen to elicit maximal interference later, in simulations where both memories are embedded to the same network. Training proceeded by simulating DC current injections to activate each cell group, with a small delay between groups allowing for STDP to strengthen connections. The awake state was followed by SWS, when the network exhibited spontaneous slow oscillations – transitions between silent Down states and active Up states (Figure 2B, right). During the awake state, recall was tested to measure performance at baseline, after training the sequence memory, and after sleep (Figure 2C). Recall was assessed by activating the first cell group and testing for pattern completion of the trained sequence (See Figure 2C insets). For single memory training (S1 or S2), recall performance was found to reliably increase for S1 (Figure 2C, top) or S2 (bottom) following both training and subsequent sleep. The coloring for the performance bars indicated which memories the network is capable of recalling: gray – neither; red – S1 only; blue – S2 only; purple – both.

**Fig. 2.**
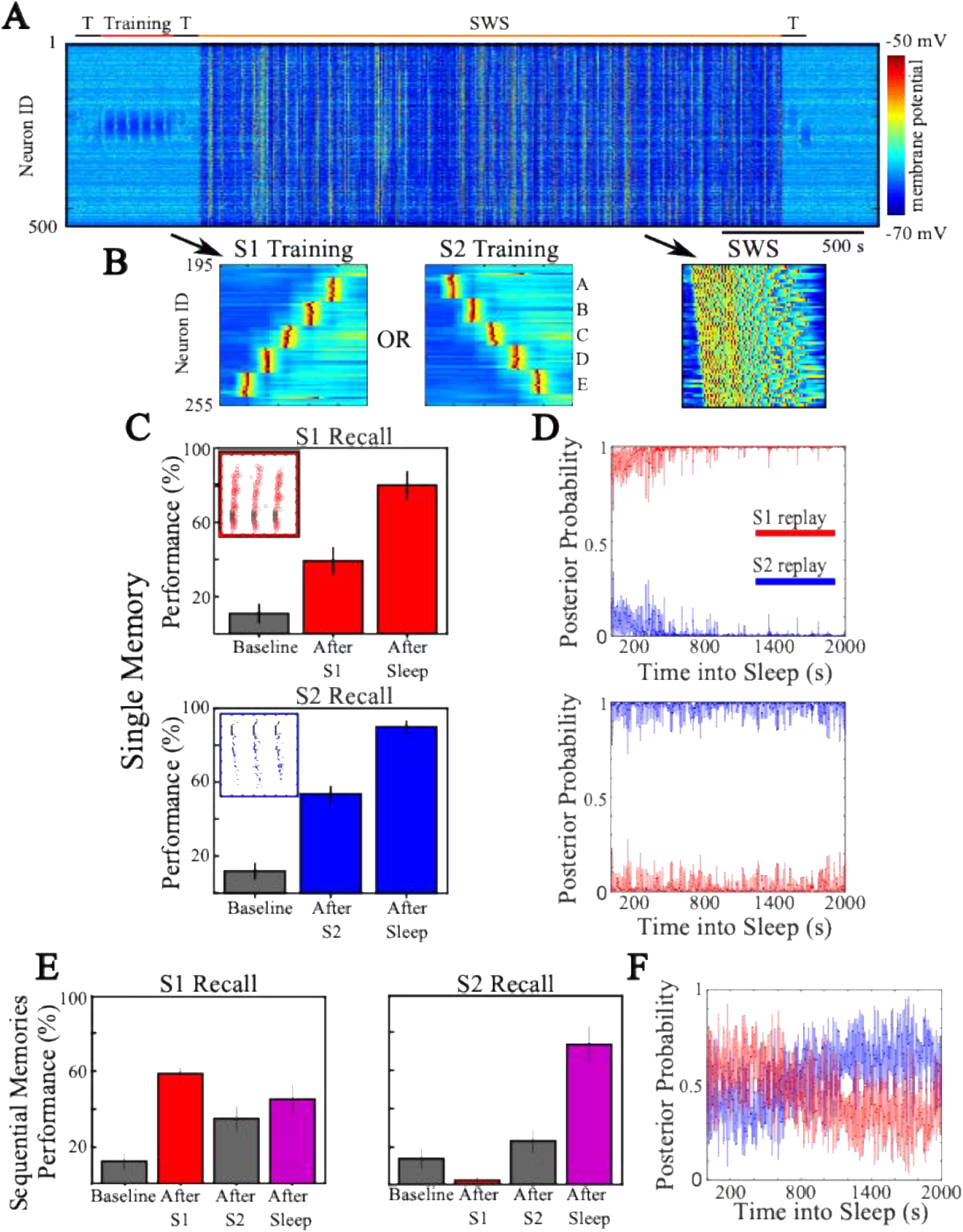
Sleep rescues interference induced by sequential training. (**A**) Network activity during an example simulation depicting all 500 PY neurons on the y-axis and the membrane potential (color scale) of each neuron over time (x-axis). The network undergoes testing periods (T; black) at baseline, after training (red), and after slow-wave sleep (SWS; orange). (**B**) Examples of network activity during one bout of S1 (left) or S2 (middle) training, and during a single Up state of SWS following a Down-to-Up transition (right). (**C**) Recall performance for S1 (top) or S2 (bottom) during all testing periods. Insets include spike rasters showing examples of pattern completion (red/blue dots) following cued recall (black dots). Bars are colored according to which memories the network can successfully pattern complete at that point in the simulation: none (gray), S1 (red), S2 (blue), or both S1 and S2 (purple). (**D**) Average probability of replaying S1 (red) or S2 (blue) during an Up state at a given time point in sleep; the top panel shows simulations in which S1 was trained, and the bottom panel shows simulations in which S2 was trained. Replay probabilities were computed using an SVM classifier trained on held-out trials from simulations in which either S1 or S2 was trained prior to sleep. (**E**) Same as (C), but for a simulation in which S2 was trained sequentially after S1 and prior to sleep; S1 recall (left) and S2 recall (right). Note the high joint recall performance (purple bar) after sleep. (**F**) Same as (D), but for the sequential-training simulation.

To evaluate differential neural reactivation, we applied PCA to reduce the dimensionality of firing-rate data during Up states in each training condition (see Supp. Figure 1A; see Methods). The Up-state trajectories following training of S1 (red) or S2 (blue) are separable in the first two principal components, indicating robust, reliable differential reactivation. We used this separation to train a linear SVM (one trial per sequence as training data; see Methods) to predict the probability that each Up state reflected an S1 versus S2 replay. Figure 2D shows these replay probabilities for each memory across SWS Up states, averaged over trials. When S1 was trained during wake, S1 replay dominated (Figure 2D, top), and the converse occurred when S2 was trained (Figure 2D, bottom). Replay probabilities varied across individual Up states, but the majority of Up states replayed the memory trained before sleep, and this bias increased over the course of sleep (see Supp. Figure 2A for single-trial replay probabilities). In sum, when a single memory was trained, sleep reliable reactivate that memory leading to post sleep increase in performance.

### Sleep can rescue interference effects induced by sequentially training memories

Next, we simulated two memories trained sequentially (S1 → S2) followed by a period of sleep. Under this paradigm, as we previously reported (Gonzalez et al., 2020), the model exhibited retroactive interference on S1 recall (Fig. 2E, left): S1 performance decreased after S2 training. We also observed prospective interference on S2 recall (Fig. 2E, right), with S2 baseline recall reduced by prior S1 training. Importantly, sleep after S1→S2 training rescued S1 from the damage caused by S2 and further improved S2 performance (first reported in (Gonzalez et al., 2020)). Figure 2F shows that the average replay probability for each sequence during sleep oscillated around 0.5, indicating both memories were replayed approximately evenly. This is well reflected by the average firing rate trajectory of the population during Up states (see Supp. Figure 1A; purple trajectory), which fell directly between the single memory trajectories. Importantly, though, for all trials individual Up states were almost always robustly classified as S1 or S2 (Supp. Fig. 2B), suggesting that different Up states were dominated by distinct memory replays in this case.

### Sleep moves the network towards stable attractors and interference can be characterized by the network escaping “memory attractor” in synaptic weight space

Next, we analyzed training- and sleep-induced model dynamics in synaptic-weight space. From the full weight space we constructed a hyperplane defined by three points - the initial weight state (O), the weight state *U*_1_ after training sequence S1, and the weight state *U*_2_ after training S2 - and evaluated recall performance for each sequence (S1, S2) at every grid point on that hyperplane. This was done in the full synaptic weights space. We then projected the system trajectories (synaptic-weight evolution during training and sleep) and this hyperplane itself into PC space (Fig. 3) and we further assessed S1/S2 performance along those trajectories. Recall-performance analysis produced contour maps on the hyperplane - the “synaptic-performance landscape” - showing S1 performance (red) and S2 performance (blue) independently (Fig. 3) and joint S1&S2 performance (purple; Supp Fig. 3), which is the average performance on the two memorizes with the absolute difference between the scores subtracted off to penalize asymmetry in performance. See Methods for PCA and landscape details.

**Fig. 3.**
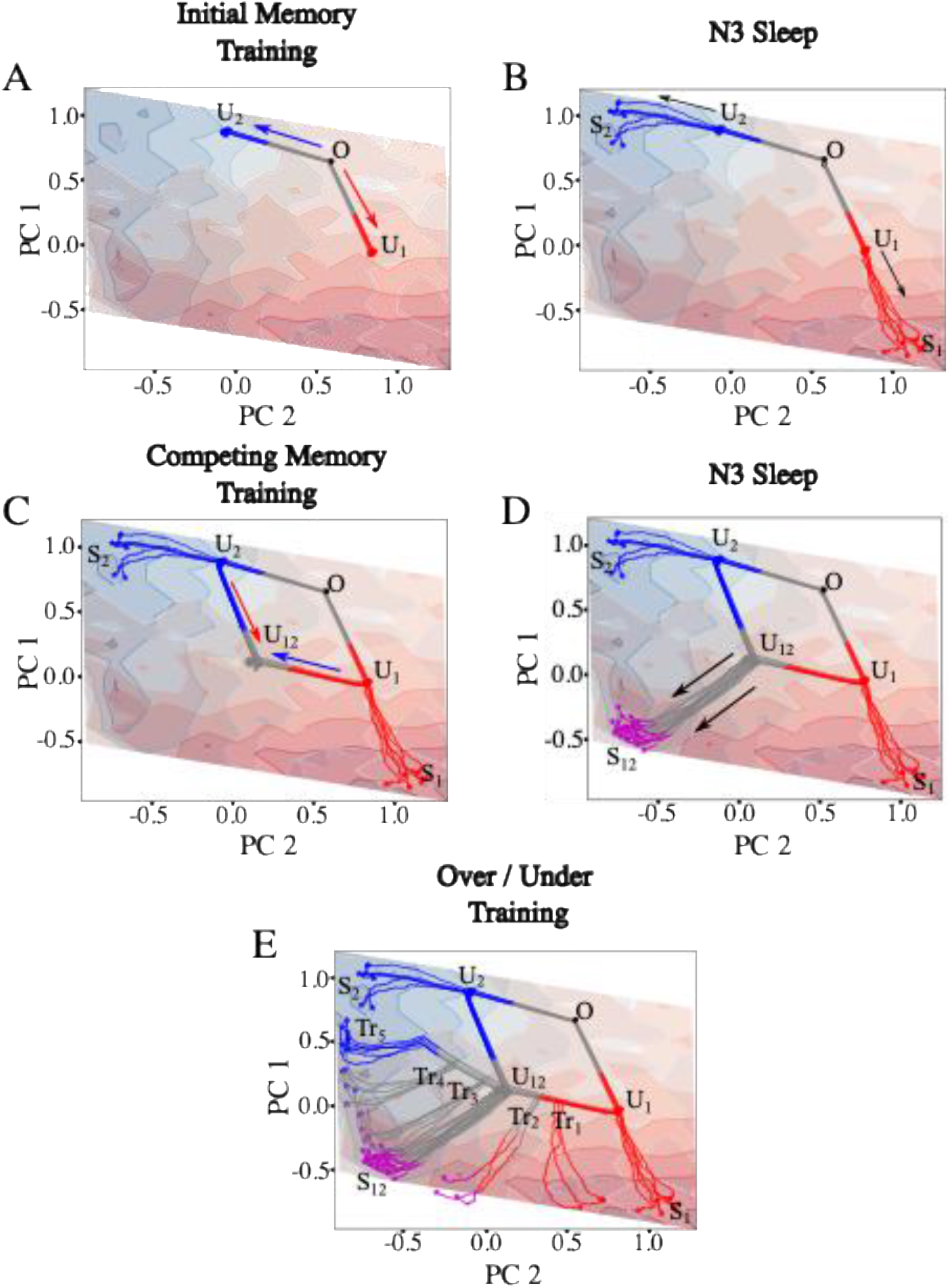
Synaptic performance landscape reveals multi-stability and fine-tuning. All panels show the evolution of the network through a dimensionality-reduced synaptic-weight space. Contour lines and color shading correspond to single-memory recall performance for S1 (red) and S2 (blue). Trajectories are colored according to which memories the network can successfully pattern complete at that point in the simulation: none (gray), S1 (red), S2 (blue), or both S1 and S2 (purple). (A) Evolution during S1 (red arrow) or S2 (blue arrow) training brings the system to synaptic states (U_1_ or U_2_) that support the corresponding memory, S1 or S2. (B) Slow-wave sleep (SWS) moves each network further along its current trajectory toward putative single-memory synaptic attractor states (S_1_ and S_2_). **(C)** Sequential training of the competing memory moves the network toward a central gray region (U_12_) where neither memory can be reliably recalled. **(D)** Subsequent SWS moves the network toward a putative joint-memory synaptic attractor (purple trajectories; S_12_), where both memories can be recalled. **(E)** Examples of undertraining (Tr_1_) and overtraining (Tr_4_, Tr_5_) of S2 following S1 training reveal the necessity of fine-tuning training durations for sleep to guide the network toward the joint-memory synaptic attractor.

Figure 3A shows that initial training of S1 (red arrow) or S2 (blue arrow) pushed the network from its initial location (O) to a new state (*U*_1_ or *U*_2_) in synaptic-weight space, leading to greater recall performance for the corresponding memory. The coloring of the trajectories reflects which memory can be robustly recalled: gray - neither; red - S1 only; blue - S2 only; purple - both. These states (*U*_1_ or *U*_2_), while delivering significant memory performance, are not stable attractors with respect to the sleep dynamics. Indeed, subsequent sleep pushes the network further along the same direction as during training, arriving at new synaptic-weight states (*S*_1_ or *S*_2_) with even greater single memory recall performance (Fig. 3B); these states behave as stable attractors under SWS dynamics, with additional sleep leading to small random fluctuations around each.

By contrast, if competing-memory training follows *U*_1_ or *U*_2_ instead of sleep (Fig. 3C), the system is driven in the opposite direction in weight space into a “region of ignorance” (gray) where neither sequence can be recalled. This low-performance state (*U*_12_) is nevertheless distinct from the initial state (O), because synaptic weights still preserve traces of both memories. Again *U*_12_ is not an attractor; consequently, subsequent sleep (Fig. 3D) moves the network into a region with high recall for both memories - the joint-memory attractor (*S*_12_). Importantly, in all cases sleep reveals stable attractor dynamics: the terminal states reached after sufficient sleep (*S*_1_, *S*_2_, *S*_12_) are stable, such that further sleep produces negligible movement around these attractors.

### Memory attractors have finite basins in synaptic weights space

Next, we tested basins of attraction of these states (*S*_1_, *S*_2_, *S*_12_) in respect to the sleep induced dynamics. Figure 3E shows examples of under/overtraining of S2 (blue) memory following initial S1 (red) training. The first undertraining example (Tr_1_) halted S2 training before the network left the basin of attraction of *S*_1_ memory attractor, and subsequent sleep pushed the network back towards the *S*_1_ attractor. Supp. Figure 4A shows the average replay probability for this case, with each memory initially replaying at roughly equal probability before S1 replay comes to dominate by the end of sleep. The second undertraining example (Tr_2_) halted S2 training just after the network left the basin of attraction of *S*_1_ attractor. In this case, subsequent sleep pushed the network towards the joint *S*_12_ memory attractor. Supp. Figure 4B shows that replay initially becomes transiently biased towards S1 before slowly returning towards replay of both memories. Similarly, the first overtraining example (Tr_3_) halts the network just after it passes the intersection in the region of ignorance, and subsequent sleep pushed it towards the joint *S*_12_ memory attractor (Supp. Figure 4C). Finally, in the last two overtraining examples (Tr_4_ and Tr_5_), the network was trained on S2 until it reached the basin of attraction of *S*_2_ memory attractor, where subsequent sleep pushed the network towards *S*_2_. Here the average replay probabilities are first balanced before S2 comes to dominate by the end of sleep (Supp. Figure 4D).

**Fig. 4.**
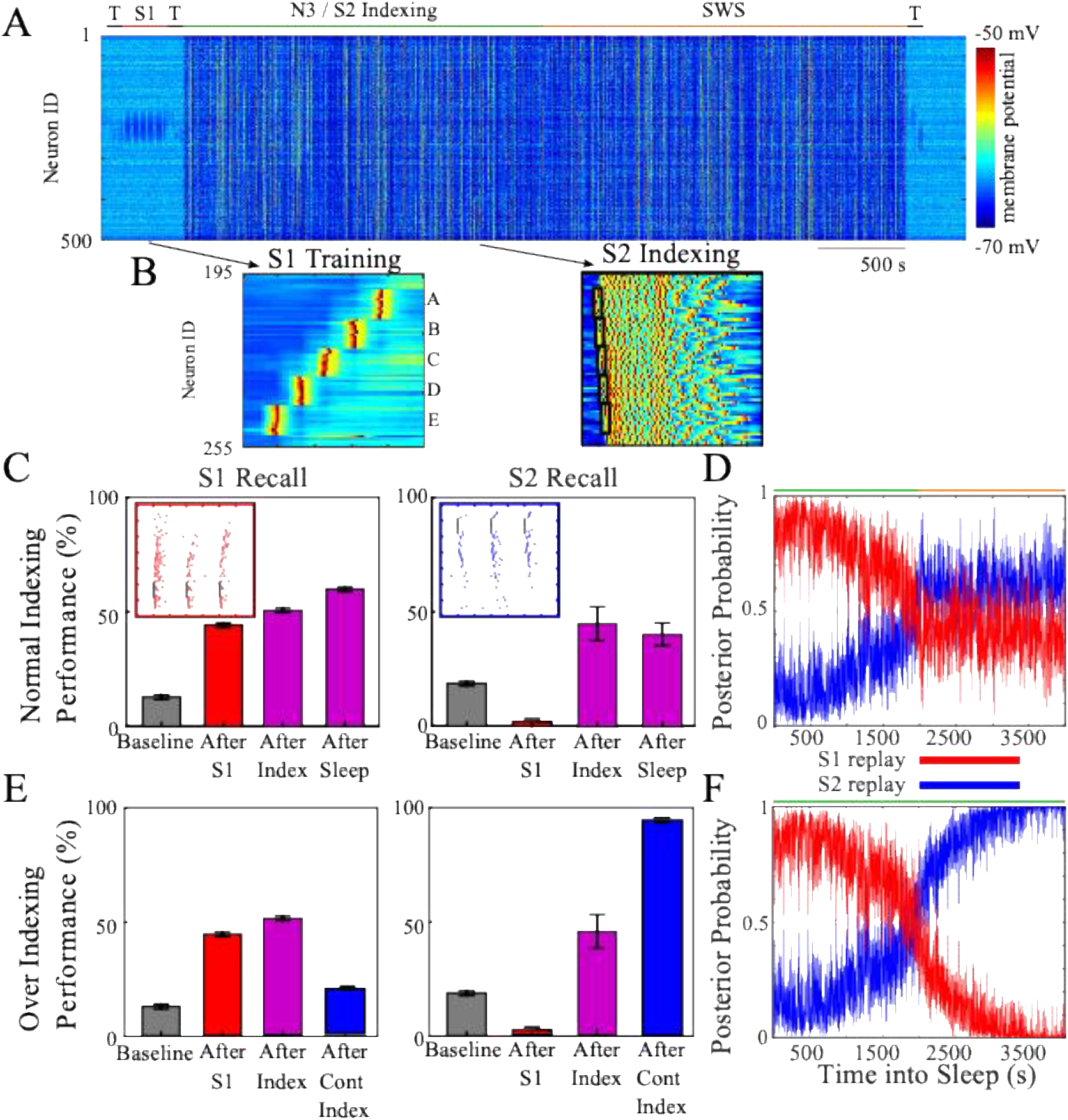
Hippocampal indexing during sleep induces consolidation without interference. (**A**) Network activity during a simulation in which the network undergoes periods of testing (T; black), training on S1 (red), slow-wave sleep (SWS) with S2 indexing (green), and SWS without indexing (orange). (**B**) Examples of network activity during one bout of S1 training (left) and during a single Up state of SWS (right), in which hippocampal indexing is simulated at the beginning of the Up state (black boxes). (**C**) Recall performance for S1 (left) and S2 (right) shows that the network consolidates S2 without interference to S1. (**D**) Average replay probability shows that S2 replay probability gradually increases during the course of indexing (green) until it becomes approximately equal to S1 replay probability by the time sleep without indexing begins (orange). (**E**) Same as (C), but for simulations in which indexing continued for the entire duration of sleep; in this case, only S2 could be recalled at the end of the simulation. (**F**) Same as (D), but for simulations in which indexing continued for the entire duration of sleep; S2 replay probability increases to unity with continual indexing.

Synaptic deletion represents a perturbation of the system in synaptic-weight space. Can sleep recover memory states disrupted by random synaptic deletion? Biological brains preserve memories over a lifetime despite constant synapse turnover, and sleep may contribute to that resilience. To test this, we started from a model trained on S1 and consolidated in sleep, then reset a fraction of the model’s synapses back to their initial (naive) values. Memory loss was gradual, and the memory was relatively resilient to synaptic damage, e.g., when 40% of synapses were reset, performance fell from ∼90% to ∼65% (Supplementary Fig. 5A). Subsequent sleep recovered a large portion of the lost accuracy: performance rose to ∼80% after a 40% synapse reset. Plotting sleep trajectories in synaptic-weight space showed the system moved back toward the attractor representing the damaged memory but settled in a nearby, slightly different state (Supplementary Fig. 5B,C), consistent with the idea that multiple weight configurations can support a given memory trace (Javed and White, 2019; Golden et al., 2022).

Taken together, these results suggest that the synaptic-weight space of the model contains at least three attractors supporting S1, S2, and the joint S1/S2 state. Furthermore, multiple attractor states represented by different synaptic configurations may support the same memory. In the simulations presented here, transitions between attractors required passing through a region of low task performance. Whether the attractor basins are fully disjoint or connected in parts of synaptic-weight space remains unclear, because it is impractical to systematically probe the model’s very high-dimensional weight space.

### Hippocampal indexing during sleep allows for new memory consolidation without interference

Although sleep could rescue retroactive and proactive interference by moving the network toward a joint memory attractor, it relied on the network passing through a region of ignorance (low task performance on both memories) to do so. The authors are unaware of any study where animals displayed the following behavioral dynamics: underwent catastrophic forgetting on the initial task without achieving significant performance improvements on the competing task but then displayed robust performance on both tasks following sleep. There are two possible solutions. First is to follow each short episode of training by sleep to rescue old memories (Golden et al., 2022). Indeed, when we interleaved short periods of S2 training with short periods of sleep, new sequence S2 was learned without damage to the old memory S1 (Supp. Figure 6). However, this approach was sensitive to the ratio and durations of sleep and S2 training phases – longer training episodes compared to sleep led to S1 forgetting, shorter training failed to learn S2. Thus, in practice this strategy would require multiple training–sleep cycles (i.e., multiple day/night cycles) and relatively low learning rates, consistent with the gradual acquisition of procedural, hippocampus-independent memories (Hong et al., 2019).

Can a new memory be trained fast but without inducing significant interference to old memories? We recently showed (Golden et al., 2025) that dual hippocampo-cortical memory system (McClelland et al., 1995; Born, 2010) provides such solution, so the hippocampal indexing during sleep allows the network to encode the competing memory with minimal interference to the initial memory. While this study (Golden et al., 2025) predicted the underlying mechanism - coordinated co-replay of old and new memories - the resulting dynamics in synaptic-weight space remain unknown.

To examine synaptic-weight dynamics during sleep indexing, we used the hippocampal indexing model introduced in (Golden et al., 2025). The simulation paradigm (Figure 4A) shows that, following a baseline test, S1 was trained (Figure 4B, left) and performance was assessed again in a post-training test. The network was then transitioned into SWS, during which hippocampal indexing of the S2 memory was triggered at each Down-to-Up transition (Isomura et al., 2006; Sanda et al., 2021). This was achieved by detecting the onset of each Up state (i.e., the Down-to-Up transition) and applying inputs to sequentially activate cell groups in the S2 order (Figure 4B, right), with a 5 ms delay between stimuli. Hippocampal indexing was restricted to the first half of the total sleep duration, as rodent studies indicate that replay of recent tasks is more robust early than late in sleep (Ji and Wilson, 2007), and declarative memory consolidation is thought to be more strongly associated with early sleep (Plihal and Born, 1999; Mednick et al., 2011; Rasch and Born, 2013). Finally, network performance on both tasks was evaluated after sleep.

Figure 4C shows that indexing produced a significant increase in S2 recall (right) that persisted during subsequent sleep without indexing, while recall performance for S1 (left) continued to rise across all stages of the simulation. The average replay probabilities (Figure 4D) indicate that the network became progressively more likely to replay S2 over the course of indexing. Terminating indexing when the two memories reached roughly equal replay probability allowed both memories to continue being replayed during subsequent sleep. Individual Up states from single-network simulations were typically classified robustly as either S1 (red) or S2 (blue) replays (Supp. Figure 2C), consistent with the sequential-training/SWS condition. If, however, indexing was allowed to continue throughout the entire sleep period, catastrophic retroactive interference occurred: the old memory S1 degraded (Figure 4E, left) while the new competing memory S2 became extremely robust (Figure 4E, right). Figure 4F shows that this outcome resulted from S2 coming to dominate replay well before the end of sleep.

Supp. Figure 1B shows that average firing-rate trajectories during Up states with indexing also fell between the single-memory trajectories described earlier. When Up states from indexing and SWS were classified into three groups based on SVM-derived reactivation probability (>95% S1 = red; >95% S2 = blue; otherwise = purple), robust S1 and S2 replays were linearly separable in two-dimensional PC space, whereas less-confident states followed intermediate trajectories consistent with mixtures of both memories.

### Hippocampal indexing moves the network to its joint memory attractor without damaging old memories

Examining system dynamics in synaptic-weight space during indexing required a third principal component, in addition to the two previously sufficient for the case in which memories were trained during wake and followed by subsequent sleep (Figure 3). This fact prohibited generation of the synaptic performance landscape, as we cannot visualize three dimensional contours.

Accordingly, Figure 5 presents two different perspectives (top and bottom panels) of a three-dimensional principal-component representation of network trajectories through synaptic-weight space. Trajectory color encodes recall capability: gray - neither memory; red - S1 only; blue - S2 only; purple - both memories.

**Fig. 5.**
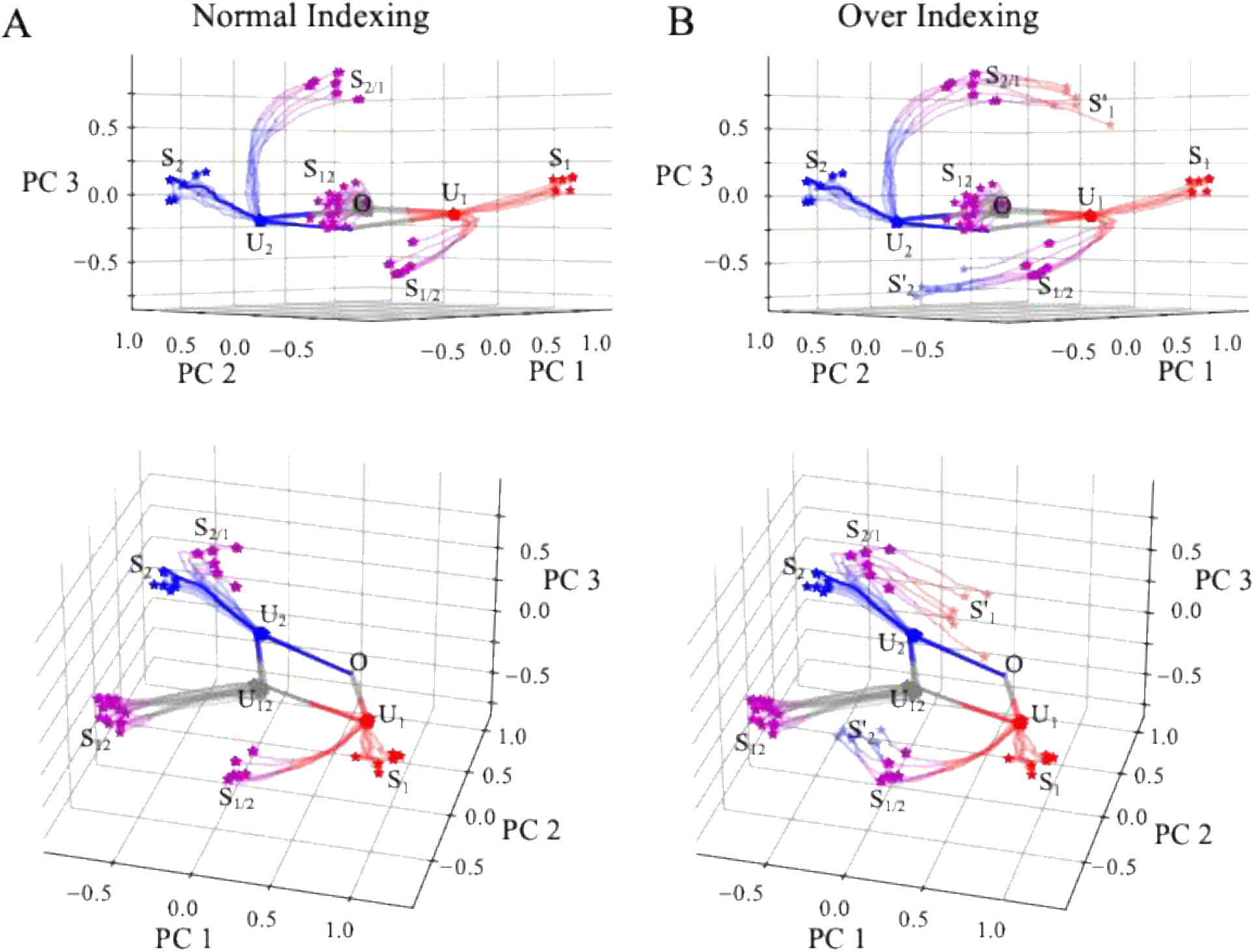
Indexing guides the network toward a joint synaptic configuration. (**A**) Evolution of the network during normal indexing (top) shows that the system moves from its initial single-memory synaptic configuration (S1, red; S2, blue), primarily along the PC3 dimension, toward a joint synaptic configuration (purple) that is stable under subsequent sleep dynamics. (**B**) Evolution during over-indexing (top) shows that if indexing is not terminated, continued hippocampal drive pushes the network away from the joint configuration and back toward a single-memory synaptic configuration corresponding to the indexed memory. The bottom panels show the same trajectories as the top panels, but with the plot rotated by 45 degrees about the PC3 axis.

As previously, naive network (O) was initially trained on S1 (or S2), moving it to the *U*_1_ (or *U*_2_) state in synaptic-weight space (Fig. 5). Subsequent indexing of the alternative memory S2 (or S1) moved the system negatively (respectively, positively) long the third principal-component dimension toward a newly identified stable attractor *S*_2/1_ (or *S*_1/2_) (Fig. 5A). In the case of over-indexing (Fig. 5B), the network was driven beyond these stable regions and continued toward the opposite single-memory attractor 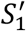 (or 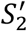). Notably, these new single-memory attractors (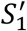 and 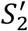) were different from original ones (*S*_1_ and *S*_2_), indicating the coexistence of multiple memory attractors in synaptic-weight space, as predicted previously (Golden et al., 2022).

Importantly, under both indexing and over-indexing, the network never traversed a “region of ignorance” and always retained robust recall of at least one memory.

### Hippocampal indexing results in sparser stable solutions

Both sequential training of two memories (S1 → S2) in wake followed by sleep (Fig. 3) and training one memory during wake followed by indexing of another during sleep (Fig. 5) produced synaptic-weight states that supported both memories (*S*_12_, *S*_2/1_, *S*_1/2_; see “purple” attractors in Figure 5). What differences exist between these states beyond their specific weight configurations? Figures 6A-B plot the PY→PY synaptic weights for all pairs of bi-directionally connected neurons for the sequential-training case (Fig. 6A) and the indexing case (Fig. 6B). We focus on bi-directional connections because unidirectional pairs (i.e., A→B but not B→A) are not subject to competition under our STDP rules. In Figs. 6A–B, the colored corner regions mark pairs in which one synapse had been prioritized for S1 (red) or S2 (blue), effectively disconnecting the opposing synapse by driving its weight to zero.

**Fig. 6.**
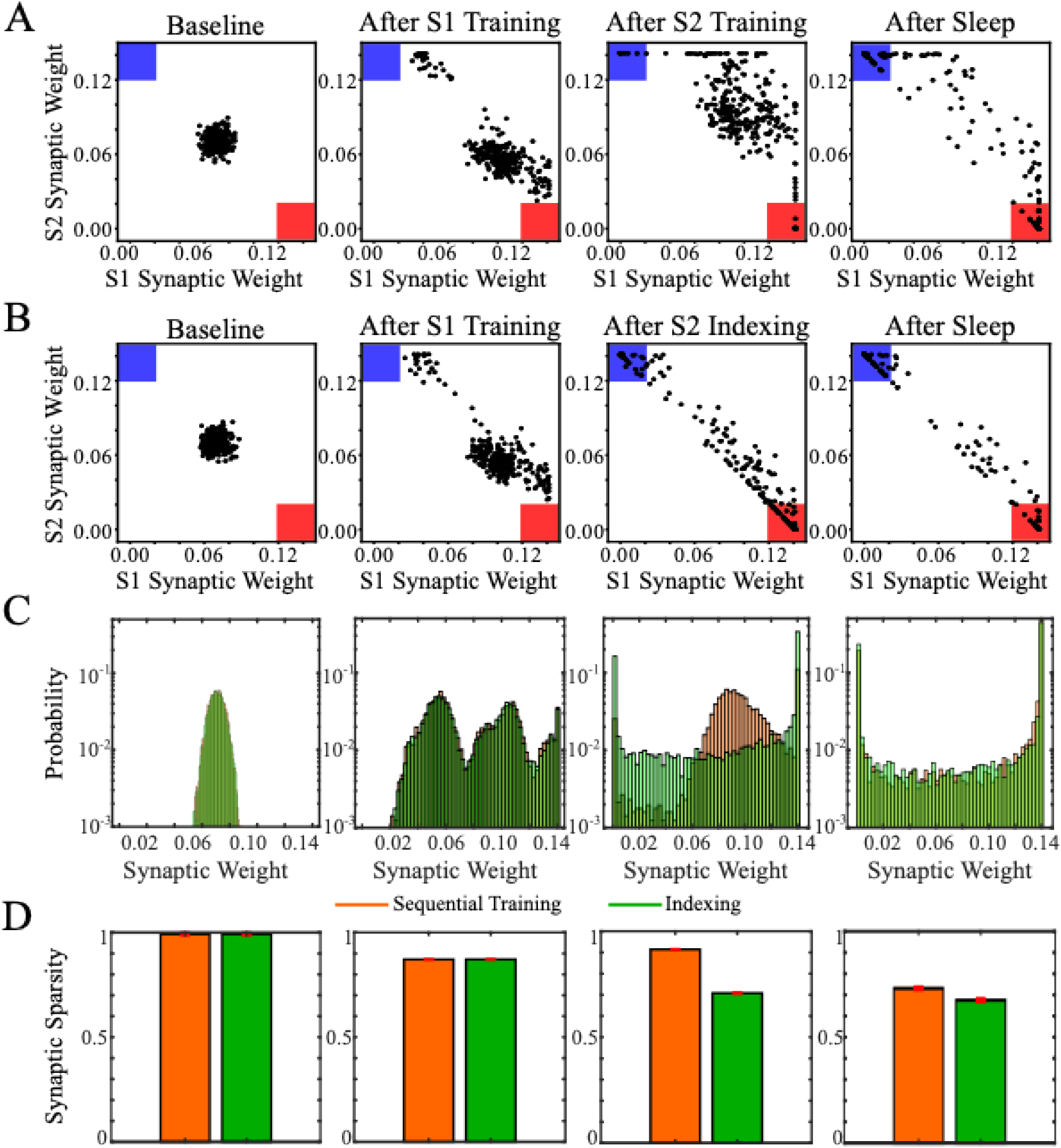
Indexing leads to sparser memory representations than sequential training and sleep. (**A–B**) Each panel shows scatter plots of synaptic weights between pairs of PY neurons in the trained region that are connected bidirectionally. The synaptic weight in the S1 direction is plotted on the x-axis, while the corresponding synaptic weight in the S2 direction is plotted on the y-axis. Colored corner regions indicate synaptic pairs that have been strongly biased toward S1 (red) or S2 (blue) at the expense of the opposing memory. (**A**) Snapshots taken during a simulation with sequential training followed by sleep. (**B**) Snapshots taken during a simulation with normal hippocampal indexing during sleep. (**C**) Distributions of synaptic weights at each time point from (A–B) for sequential training (orange) and normal indexing (green), plotted with a logarithmic y-axis. (**D**) Average synaptic density of the weight matrix across simulations with sequential training (orange) and normal indexing (green). Error bars indicate standard deviation.

In both paradigms the initial weight values were roughly Gaussian (Figs. 6A–B, left), and the bulk of the density shifted toward the red corner after S1 training (Figs. 6A–B, middle-left). Beyond this point, the two paradigms diverged. Both sequential training during wake and indexing during sleep produced more symmetric density distributions overall (Fig. 6A-B, middle-right), indicating a roughly equal allocation of synaptic resources to the two memories. However, S2 training during wake (Fig. 6A, middle-right) pushed the density much farther into the top-right corner than S2 indexing during sleep (Fig. 6B, middle-right), which kept the density localized around the diagonal connecting the red and blue corners (Y = −X). This pattern implies that sequential training during wake yields many bi-directional pairs that are strong in both directions - an ambiguity that prevents those pairs from contributing to differential memory encoding. By contrast, hippocampal indexing preferentially segregates (orthogonalizes) synaptic resources between the two memories. In both paradigms, subsequent sleep without indexing amplified any preexisting S1/S2 bias in each bi-directional pair, pushing pairs further into either the red or blue corner (Figs. 6A–B, right). However, sleep following wake training required much larger synaptic changes to correct weight saturation.

Figure 6C shows the synaptic-weight distributions obtained after the sequential-training during wake (orange) and indexing during sleep (green) paradigms for the corresponding simulation phases shown above. The distribution reached by indexing during sleep is noticeably sparser than the distribution produced by sequential training (Fig. 6C, middle-right), consistent with greater segregation of synaptic resources under indexing.

To quantify the density of learned synaptic representations, we used a measure originally developed to compute the sparseness of neural population codes (Treves and Rolls, 1994) (see **Methods** for more details). Low values indicate that information is carried by a small number of synapses (low-density synaptic representation), whereas values near 1 indicate that information is broadly distributed across synapses (high-density representation). Figure 6D plots this density at the same four time points shown in Fig. 6A–C. In both paradigms, initial S1 training reduced representational density. However, sequential S2 training (orange bars) increased density, while S2 indexing during SWS (green bars) further reduced it - a pattern consistent with stronger interference following sequential training during wake than with sleep indexing. Finally, subsequent sleep in both paradigms further decreased density, but sleep indexing led to significantly sparser - and therefore more resource-efficient - synaptic-weight configurations than sequential training during wake.

## DISCUSSION

The human brain contains an estimated 10^14^ synapses and is in constant interaction with the environment, leading to continual perturbations of its synaptic weight matrix. As demonstrated in artificial neural networks (ANNs), continuous acquisition of new information can induce interference and catastrophic forgetting (Mccloskey and Cohen, 1989; McClelland et al., 1995; French, 1999; Hayes et al., 2021), and can also drive synaptic saturation and loss of plasticity (Dohare et al., 2024). How, then, does the vast pool of memories accumulated over a lifetime persist despite ongoing perturbations, and without requiring explicit relearning?

The critical role of sleep in learning and memory is supported by a vast, interdisciplinary literature spanning psychology and neuroscience (Paller and Voss, 2004; Walker and Stickgold, 2004; Oudiette et al., 2013; Rasch and Born, 2013; Stickgold, 2013). The central idea is that sleep contributes to memory consolidation through repeated reactivation, or replay, of memory traces acquired during awake learning (Hennevin et al., 1995; Paller and Voss, 2004; Mednick et al., 2011; Oudiette et al., 2013; Rasch and Born, 2013; Lewis et al., 2018; Wei et al., 2018). However, how sleep reshapes synaptic representations of old and new memories in high-dimensional synaptic weight space - so that new knowledge is embedded without damage to old memories - remains unknown.

Previous modeling studies showed that sleep-dependent replay protects previously learned memories through structured reactivation of old and new traces within cortical circuits (Gonzalez et al., 2020; Golden et al., 2025). Gonzalez et al. (2020) demonstrated that sleep reshapes the synaptic footprints of coexisting cortical memories under competition for shared neuronal and synaptic resources, while Golden et al. (2025) introduced Structured Cortical Replay (SCoRe), in which slow-wave sleep interleaves replay of familiar cortical memories with newly acquired hippocampus-dependent traces to support continual learning. However, these studies did not characterize how sleep-driven plasticity modifies synaptic-weight space to give rise to stable synaptic configurations. Here, we address this gap by identifying sleep-driven synaptic attractors - synaptic configurations that are stable with respect to autonomous sleep dynamics - and by showing how hippocampo–cortical interactions move the system between these attractors to enable continual learning.

From a dynamical-systems perspective, brain networks comprise variables evolving on distinct time scales: fast variables, such as neuronal activity, and slow variables, such as synaptic weights. When synaptic weights are assumed to be fixed, network dynamics are determined by the resulting synaptic-weight landscape, in which each memory corresponds to an attractor of the neuronal activation dynamics. Starting from random activity, the network converges to the active state represented by one such attractor, as proposed in classic models like the Hopfield network (Hopfield, 1982), which have been generalized to include temporal sequence dynamics (Kleinfeld, 1986; Hopfield, 1996; Chaudhry et al., 2023). Learning then modifies synaptic weights, reshaping this landscape and thereby governing fast dynamics, for example by creating new memory attractors. The assumption of fixed synaptic connectivity is justified because synaptogenesis typically occurs on a much slower time scale than neuronal activity.

However, this classical picture neglects another fundamental component of biological systems: sleep. During sleep, sensory input is largely absent, yet internal dynamics actively reshape the synaptic landscape. Although synaptic changes remain slow, they are substantial on the time scale of a typical 8-hour sleep period in humans that recurs daily. As a result, the synaptic landscape governing fast activation dynamics differs before and after sleep. Understanding how sleep reshapes this landscape is therefore essential both for understanding biological sleep and for developing continual-learning artificial systems.

We propose that synaptic configurations supporting strong memories are attractors in synaptic-weight space. Sleep does not randomly perturb synaptic weights; instead, by replaying each stored memory, it deepens the attractor basin around that memory’s configuration. The net effect is that the overall weight configuration - the one jointly encoding all memories - is not disrupted by sleep but reinforced by it.

It is important to distinguish two kinds of attractors. *Memory attractors* are attractors in neuronal activity space: each encodes a single memory, and convergence to one corresponds to retrieving that memory. *Synaptic attractors* are attractors in synaptic-weight space: each represents a stable network configuration — encoding none, one, or multiple memories — and convergence to one corresponds to the network settling into a weight configuration that supports a particular set of memories. Crucially, individual memories correspond to attractors only in neuronal activity space; attractors in synaptic-weight space are higher-order objects that determine *which* memory attractors exist. We use this terminology — synaptic attractors versus memory attractors — throughout.

Both daytime learning and sleep reshape the synaptic landscape. Strong perturbations - e.g., learning a new task - may push the system toward the basin of attraction of a synaptic state that supports only the new memory (overtraining), inducing forgetting, or toward an attractor state that supports both memories (a joint state), enabling continual incremental learning. While daytime experiences modify synaptic weights, forming new memory attractors and, as a result, damaging old ones, sleep-induced replay tends to modify the synaptic landscape to return the system toward synaptic weights configurations (synaptic attractors) that support all sufficiently strong memories. Indeed, we recently successfully implemented sleep-like dynamics in feedforward and recurrent ANNs, where a sleep phase prevented catastrophic forgetting and improved generalization (Tadros et al., 2020a; Tadros et al., 2022; Delanois et al., 2023; Kubo et al., 2025).

Figure 7 summarizes these concepts with illustrative schematics. Two states (S1 and S2) represent stable attractors in synaptic weights space supporting individual memories (Memory 1 and Memory 2, respectably); the third state, S12, corresponds to a synaptic attractor that supports both memories. Curves M1 and M2 illustrate other possible synaptic configurations that support Memory 1 and Memory 2, respectively, but are not necessarily attractors under sleep dynamics (see below). When initial training brings the system into the vicinity of a single-memory attractor (e.g., S1), subsequent sleep replay modifies synaptic weights and drives convergence toward that attractor. Small perturbations away from S1 are compensated by sleep dynamics that return the system to S1. Learning a new task during wake shifts the system away from the old attractor (S1) toward the new single-memory attractor (S2) (Fig. 7A–C). With subsequent sleep, the system may transition to the joint-memory attractor (S12) (Fig. 7A), but this outcome requires a finely tuned training duration so that the system enters the basin of attraction of S12. Otherwise, the network may fail to consolidate the new memory (proactive interference; return to S1, Fig. 7B) or undergo catastrophic forgetting (retroactive interference; transition to S2, Fig. 7C). This raises the question of whether it is possible to avoid this sensitivity and achieve reliable continual learning that benefits from sleep dynamics.

**Fig. 7.**
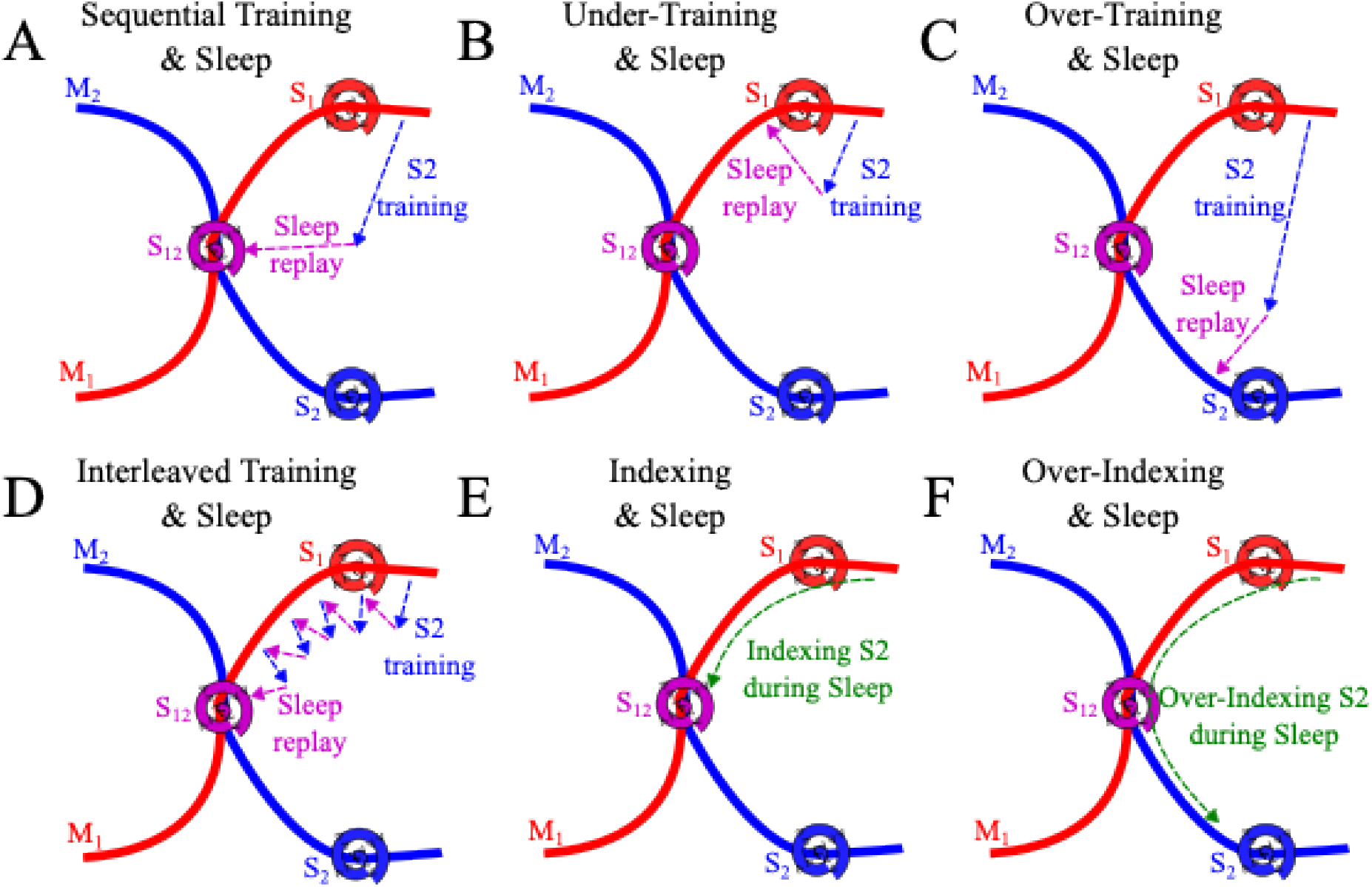
Systems consolidation prevents interference through interleaved replay within Up states. Cartoon schematics illustrating network dynamics in synaptic-weight space for (A) new (S2) memory training followed by sleep, (B) new memory under-training followed by sleep, (C) new memory over-training followed by sleep, (D) interleaving new memory training with sleep, (E) hippocampal indexing of a new memory during sleep, and (F) over-indexing of a new memory during sleep. Spirals represent synaptic configurations that appear stable and attracting under autonomous sleep dynamics, corresponding to synaptic attractor states.

Our current study, together with previous work (Golden et al., 2022), reveals that multiple synaptic weight configuration may support a given memory trace. [In (Golden et al., 2022), the full set of such configurations was described using the concept of memory manifolds; however, for clarity of presentation, we do not use this terminology here.] While only a subset of these configurations is stable with respect to autonomous sleep dynamics, all of them support a given memory (or set of memories). For example, all points along curve M1 (Fig. 7) support Memory 1, but only S1 and S12 correspond to synaptic attractors in respect to sleep induced synaptic weights dynamics, and they differ in whether additional memories are also encoded. The goal of continual learning is therefore to move the system from a synaptic attractor that supports only previously learned memories (e.g., S1) to a synaptic attractor that supports both old and new memories (e.g., S12), without passing through states that eliminate previously stored memories.

One possible solution is to interleave new training with periods of sleep to repair any degradation of previously learned tasks (Fig. 7D). For procedural, hippocampus-independent memories, new learning is typically slow, and interference with existing memories after each training session is therefore minimal. In this regime, subsequent sleep episodes can compensate for accumulated damage and return the system toward synaptic configurations that preserve older memories. As we report here and in previous work (Golden et al., 2022), this strategy can be effective but remains sensitive to the relative durations of training and sleep episodes, requiring fine-tuning of learning-period length.

We propose that a memory system involving a fast-learning hippocampus and a slow-learning cortex, as in declarative memory, provides a more robust solution (Fig. 7E). In this case, hippocampal indexing during sleep promotes coordinated replay of newly learned information together with relevant older cortical memories (Golden et al., 2025). This sleep-driven synaptic dynamics allows the system to transition from synaptic configurations supporting only old memories toward joint synaptic attractors that support multiple memories, while remaining within the set of synaptic configurations that support previously learned memories. Once the system enters the basin of attraction of such a joint state (e.g., S12), subsequent sleep without hippocampal indexing can further stabilize this configuration. Although excessive hippocampal indexing can still disrupt older cortical memories (Fig. 7F), the duration of hippocampo–cortical interaction in the brain is regulated by intrinsic hippocampal mechanisms and is largely independent of learning duration, typically being limited to the first 30–60 minutes of post-learning sleep (Wilson and McNaughton, 1994; Ji and Wilson, 2007; Eschenko et al., 2008).

Why are not all memory-supporting synaptic configurations stable under SWS dynamics? We speculate that stability during sleep requires self-consistency between replay activity and sleep-specific plasticity rules. Synaptic configurations that fail to generate replay patterns compatible with STDP during slow oscillations are not fixed points of the sleep dynamics and therefore evolve toward alternative attractor states.

It is interesting to note that within this framework, the complementary learning-systems architecture can be viewed as an efficient implementation of interleaved training. According to the Structured Cortical Replay (SCoRe) hypothesis (Golden et al., 2025) slow-wave sleep interleaves replay of familiar cortical memories with newly acquired hippocampus-dependent traces to support continual learning. From this perspective, both procedural and declarative memory systems implement interleaved learning - a long-proposed solution to catastrophic forgetting. Consistent with this view, extensive prior work has shown that interleaved training mitigates catastrophic forgetting in ANNs (McClelland et al., 1995; Hasselmo, 2017; Flesch et al., 2018) and SNNs (Evans and Stringer, 2012; Higgins et al., 2017), motivating replay-based algorithms that mix past and recent experiences during learning (reviewed in (Hayes et al., 2021)). We propose that the brain achieves the same functional outcome without explicit storage of past data, either by separating slow wake learning from spontaneous reactivation during sleep (procedural memory), or by embedding spontaneous replay of older traces within hippocampo– cortical interactions during sleep (declarative memory). In both cases, sleep dynamics push the system toward synaptic attractor states that support both old and new memories, thereby avoiding catastrophic forgetting.

In sum, our study suggests that interactions between the hippocampus and neocortex during slow-wave sleep provide optimal dynamics for guiding the system between stable synaptic attractors that preserve previously learned memories while incorporating new ones. We observed complex synaptic weight dynamics during SWS, with multiple stable synaptic attractors representing individual memories as well as joint attractor states supporting the coexistence of multiple memory traces. These synaptic configurations are stable with respect to autonomous sleep dynamics. Importantly, although hippocampal indexing drives transitions between basins of attraction toward states supporting multiple memories, previously stored memories are preserved throughout this process.

## METHODS

### Network architecture

Throughout this study, we make use of a modified version of a thalamocortical network which has been previously described in detail (Wei et al., 2016; Gonzalez et al., 2020). In brief, the network consisted of a cortical module containing 500 excitatory pyramidal neurons (PYs) and 100 inhibitory interneurons (INs), and a thalamic module containing 100 excitatory thalamocortical neurons (TCs) and 100 inhibitory reticular interneurons (REs). Connectivity in the network was determined by cell type and a local radius (see Fig. 1), and excitatory synapses were mediated by AMPA and/or NMDA currents, while inhibitory synapses were mediated by GABA_A_ and/or GABA_B_ currents.

In the cortex, PYs synapsed onto PYs and INs with a radii of R_AMPA(PY-PY)_ = 20, R_NMDA(PY-PY)_ = 5, R_AMPA(PY-IN)_ = 1, and R_NMDA(PY-IN)_ = 1. All connections were deterministic within these radii, expect for AMPA synapses between PYs, which had a 60% probability of connection.

Additionally, INs synapsed onto PYs with a radius of R_GABA-A(IN-PY)_ = 5. In the thalamus, TCs synapsed onto REs with a radius of R_AMPA(TC-RE)_ = 8 and REs synapsed onto REs and TCs with radii of R_GABA-A(RE-RE)_ = 5, R_GABA-A(RE-TC)_ = 8, and R_GABA-B(RE-TC)_ = 8. Between the cortex and thalamus, TCs synapsed onto PYs and INs with radii of R_AMPA(TC-PY)_ = 15, R_AMPA(TC-IN)_ = 3, while PYs synapsed onto TCs and REs with radii of R_AMPA(PY-TC)_ = 10, and R_AMPA(PY-RE)_ = 8.

### Wake – Sleep transitions

To model the state transitions between awake and N3 sleep, we modulated the intrinsic and synaptic currents of our neuron models to account for differing concentrations of neuromodulators that partially govern these arousal state transitions. As these mechanisms have been described in detail in (Krishnan et al., 2016), here we will simply outline the approach. The model included the effects of changing acetylcholine (ACh), histamine (HA), and GABA concentrations as follows: ACh – by modulating the potassium leak current in all cell types, as well as excitatory AMPA synapses within the cortex; HA – by modulating the hyperpolarization-activated cation current in TC cells; and GABA – by modulating inhibitory GABAergic synapses within the cortex and thalamus. To transition the network from awake to sleep, we modeled the effects of reduced ACh and HA but increased GABA concentrations to reflect experimental observations (Vanini et al., 2012).

### Intrinsic currents

All cell types were modeled using the Hodgkin-Huxley formalism, and cortical PYs and INs contained dendritic and axo-somatic compartments that have been previously described (Wei et al., 2018). The dynamics of the membrane potential were modeled according to:

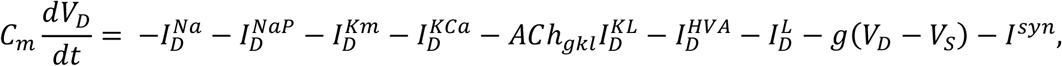

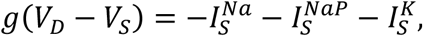

where *C*_*m*_ is the membrane capacitance, *V*_*D*_,_*S*_ are the dendritic and axo-somatic membrane voltages respectively, *I*^*Na*^ is the fast sodium (Na^+^) current, *I*^*NaP*^ is the persistent Na^+^ current,

*I*^*Km*^ is the slow voltage-dependent non-inactivating potassium (K^+^) current, *I*^*KCa*^ is the slow calcium (Ca^2+^)-dependent K^+^ current, *ACh*_*gkl*_ represents the change in K^+^ leak current *I*^*KL*^ which is dependent on the level of ACh during the different arousal states, *I*^*HVA*^ is the high-threshold Ca^2+^ current, *I*^*L*^ is the chloride (Cl^-^) leak current, *g* is the conductance between the dendritic and axo-somatic compartments, and *I*^*syn*^ is the total synaptic current input to the neuron. IN neurons contained all intrinsic currents present in PY with the exception of the *I*^*NaP*^. All intrinsic ionic currents (*I*^*j*^) were modeled in a similar form:

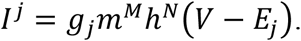

where *g*_*j*_ is the maximum conductance, *m* (activation) and *h* (inactivation) are the gating variables, *V* is the voltage of the compartment, and *E*_*j*_ is the reversal potential of the ionic current. The gating variable dynamics are described as follows:

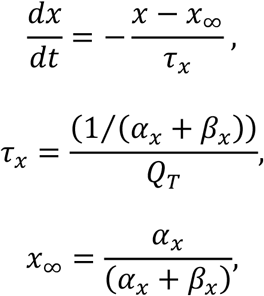

where *x* = *m* or *h, τ* is the time constant, *Q*_*T*_ is the temperature related term, *Q*_*T*_ = *Q*^((*T*−23)/10)^ = 2.9529, with *Q* = 2.3 and *T* = 36.

In the thalamus, TCs and REs contained a single compartment with membrane potential dynamics given by:

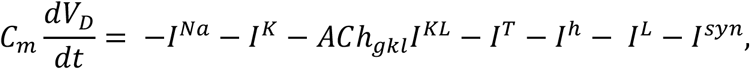

where *I*^*Na*^ is the fast Na^+^ current, *I*^*K*^ is the fast K^+^ current, *I*^*KL*^ is the K^+^ leak current, *I*^*T*^ is the low-threshold Ca^2+^ current, *I*^*h*^ is the hyperpolarization-activated mixed cation current, *I*^*L*^ is the Cl^-^ leak current, and *I*^*syn*^ is the total synaptic current input to the neurons. The *I*^*h*^ current was only expressed in TCs. The influence of histamine (HA) on *I*^*h*^ was implemented as a shift in the activation curve by *HA*_*gh*_ as described by:

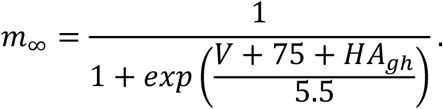

### Synaptic currents

The equations for our synaptic current models have been described in detail in our previous studies (Krishnan et al., 2016; Wei et al., 2018). To model the effects of ACh and GABA, we modified the standard equations as follows:

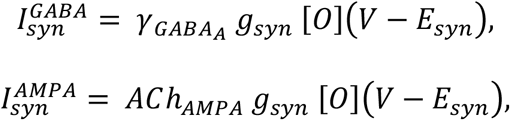

where *g*_*syn*_ is the maximal conductance at the synapse, [*O*] is the fraction of open channels, and *E*_*syn*_ is the channel reversal potential (E_GABA-A_ = -70 mV, E_AMPA_ = 0 mV, and E_NMDA_ = 0 mv). The parameter 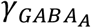 modulated the GABA synaptic currents for IN-PY, RE-RE, and RE-TC connections. For INs 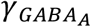 was 0.22 and 0.44 for awake and N3 sleep, respectively, while for REs 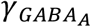 was 0.6 and 1.2. *ACh*_*AMPA*_ defined the influence of ACh levels on AMPA synaptic currents for PY-PY, TC-PY, and TC-IN. For PYs *ACh*_*AMPA*_ was 0.133 and 0.4332 for awake and N3 sleep, respectively, while for TCs *ACh*_*AMPA*_ was 0.6 and 1.2.

In addition to spike-triggered post-synaptic potentials (PSPs), spontaneous miniature PSPs (mPSPs) were implemented for both excitatory and inhibitory synapses within the cortex. The dynamics are similar to the typical PSPs described above, but the arrival times were governed by an inhomogeneous Poisson process where the next release time *t*_*release*_ is given by:

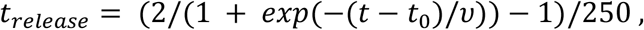

where *t*_0_ is the time of the last presynaptic spike, and *v* was the mPSP frequency 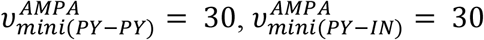 and 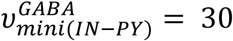. The maximum conductances for 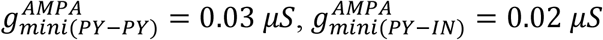 and 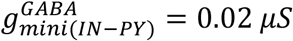.

Finally, short-term synaptic depression was also implemented in AMPA synapses within the cortex. To model this phenomenon, the maximum synaptic conductance was multiplied by a depression variable (*D* ≤ 1), which represents the amount of available “synaptic resources” as described in (Bazhenov et al., 2002). This short-term depression was modeled as follows:

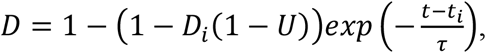

where *D*_*i*_ is the value of *D* immediately before the *i*_*th*_ event, (*t* − *t*_*i*_) is the time after the *i* _*th*_ event, *U* = 0.073 is the fraction of synaptic resources used per action potential, and *τ* = 700*ms* is time constant of recovery of synaptic resources.

### Spike-timing-dependent plasticity

The potentiation and depression of AMPA synapses between PYs were governed by the following spike-timing-dependent plasticity (STDP) rule:

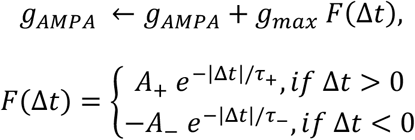

where *g*_*max*_ was the maximal conductance of *g*_*AMPA*_, *F* was the STDP kernel, and □*t* was the relative timing of the pre- and post-synaptic spikes. The maximum potentiation/depression were set to A_+/-_ = 0.002, while the time constants were set to t_+/-_ = 20 ms. A_-_ was reduced to 0.001 during training to reflect the effects of changes in acetylcholine concentration during focused attention on synaptic depression during task learning observed experimentally (Blokland, 1995; Shinoe et al., 2005; Sugisaki et al., 2016).

### Sequence training and testing

Training and testing of memory sequences was performed similarly to our previous study (Wei et al., 2018). In brief, each sequence was comprised of the same 5 groups of 10 PYs (i.e PYs 200 - 249), with Sequence 1 (S1) ordered E(240-249), D(230-239), C(220-229), B(210-119), A(200-209), and Sequence 2 (S2) ordered A(200-209), B(210-219), C(220-229), D(230-239), E(240-249). Each training bout consisted of sequentially activating each group via a 10 ms direct current pulse with a 5 ms delay between group activations. Training bouts occurred every 1 s during the training period. This training structure was chosen to ensure strong interference between S1 and S2 according to our STDP rule. Test bouts occurred every 1 ms during testing periods, in which only the first group in each sequence was activated (E for S1; A for S2), and recall performance was measured based on the extent of pattern completion for the remainder of the sequence within a 350 ms window.

### Data Analysis

All analyses were performed with standard MatLab and Python functions. Data are presented as mean □ standard error of the mean (SEM) unless otherwise stated. For each experiment a total of 6 simulations with different random seeds were used for statistical analysis.

### Sequence performance measure

A detailed description of the performance measure used during testing can be found in (Wei et al., 2018) and the code is available in (https://github.com/o2gonzalez/sequencePerformanceAnalysis) (Gonzalez et al., 2020). Briefly, the performance of the network on recalling a given sequence following activation of the first group of that sequence was measured by the percent of successful sequence recalls. We first detected all spikes within the predefined 350 ms time window for all 5 groups of neurons in a sequence. The firing rate of each group was then smoothed by convolving the average instantaneous firing rate of the group’s 10 neurons with a Gaussian kernel with window size of 50 ms. We then sorted the peaks of the smoothed firing rates during the 350 ms window to determine the ordering of group activations. Next, we applied a string match (SM) method to determine the similarity between the detected sequences and an ideal sequence (ie. A-B-C-D-E for S1). SM was calculated using the following equation:

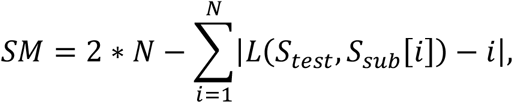

where *N* is the sequence length of *S*_*test*_, *S*_*test*_ is the test sequence generated by the network during testing, *S*_*sub*_ is a subset of the ideal sequence that only contains the same elements of *S*_*test*_, and *L*(*S*_*test*_, *S*_*sub*_[*i*]) is the location of the element *S*_*sub*_[*i*] in sequence *S*_*test*_. *SM* was then normalized by double the length of the ideal sequence. Finally, the performance was calculated as the percent of recalled sequences with *SM* ≥ *Th* = 0.8, where *Th* is a threshold indicating that the recalled sequence must be at least 80% similar to the ideal sequence to be counted as a successful recall as previously done in (Wei et al., 2018).

### Representation of firing rate space

To visualize dimensionality reduced trajectories during Up-states in firing rate space, we took single random seeds of a simulations in which only S1 or only S2 was trained prior to N3 sleep and detected Up-states in each. We then generated spike rasters of the PY activity in the trained region during each Up-state, and converted these into firing rates by taking a moving average of the spike rasters with a sliding window length of 10 ms, resulting in a set of *N-*by-*T*_*j*_ matrices, where *N* = 50 was the number of PYs, and *T*_*j*_ was the duration of *j*^*th*^ Up-state with *j* = 1, … *N*_*up*_. The firing rate matrices were then interpolated to be the same duration across each up state, and concatenated, resulting in an *N*-by-(*T*_max_\**N*_*up*_) matrix where *T*_*max*_ was the duration of longest Up-state across all data sets. Principal components analysis (PCA) was then performed on the firing rate data to reduce it from 50 to 2 dimensions. This linear PCA kernel was then applied to the data from all random seeds for a particular simulation paradigm, and the mean and standard deviation of PCs 1 and 2 were plotted for visualization.

### Replay probability

To estimate the probability of S1 and S2 being replayed during a given Up-state, we took the interpolated firing rate data for each Up-state that was used to train the PCA kernel, unrolled the *N*-by-*T*_*j*_ matrices into (*N*x*T*_*j*_)-dimensional column vectors, providing the observations to train a linear support vector machine (SVM), with labels determined by whether the Up-state came from the simulation with S1 or S2 training before N3 sleep, and scores were transformed to posterior probabilities. This SVM was then used to predict the posterior probabilities of S1 and S2 replay for each Up-state for a given random seed of a particular simulation paradigm. To compute the average posterior probabilities, we first interpolated the data so that each random seed had the same number of data points – specifically, the maximum number of Up-states in a single simulation from that paradigm.

### Representation of synaptic weight space

In order to visualize the trajectories of the network through synaptic weight space, we trained a linear PCA kernel on the synaptic weight timeseries data of all synapses in the trained region from every random seed of each simulation paradigm discussed in the paper, except for the simulations involving over/under-training, or synaptic resetting. The data was then transformed into PC space, with either 2 or 3 dimensions retained for plotting.

### Performance contours in synaptic weight space

In order to construct the recall performance contours, we first estimated the locations of the potential attractor sites by computing the centroids of the final weight state configurations for the S1 training, S2 training, and the sequential training simulation paradigms in full-dimensional weight space, found the unique 2-dimensional planar subspace which these three points define, and then densely sampled weight state configurations from this subspace. These weight state configurations were then input into the network model to have recall performance evaluated. These sampled weight states were then projected into 2-dimensional PC space along with the corresponding mean performance values for S1 and S2 individually (Figure 3; red and blue), and S1&S2 jointly (Supp Figure 3; purple), and a mesh with contour gradients was computed for each set of recall performance measures.

### Desnity of synaptic weight states

In order to compute the density of synaptic weight matrices, we adpated a measure previously conceived to compute the sparsity of neural population codes, operating over real-valued firing rates (Treves and Rolls, 1994),

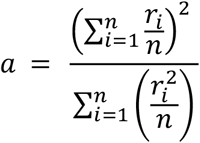

where *a* is the density of the population code, *r*_*i*_ is the firing rate of the *i*th neuron, and *n* is the total number of neurons. The density here is maximal at 1 when all neurons have the same firing rate, 1/*n* when only one neuron responds, and zero when the population is silent.

To adapt this to compute the density of synaptic weight states, we first unrolled the synaptic weight matrix into a vector and then removed all non-existent synapses (i.e. zero-valued) from the vector so that we only computed over the number of existing synapses rather than potential synapses. We then computed

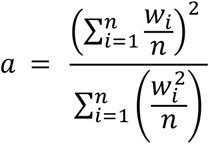

where *a* is the density of the weight state, *w*_*i*_ is the firing rate of the *i*th synapse, and *n* is the total number of synapses. Thus, the density here is maximal at 1 when all synapses have the same strength, and near 1/*n* when only one synapse is above its minimum strength value (note that existing synapses could not drop to zero strength).

## ACKNOWLEDGEMENTS

This work was supported by NIH (1R01MH125557, 1RF1NS132913, RF1NS132041), NSF (2223839).

## FIGURES

**Supp. Fig. 1.**
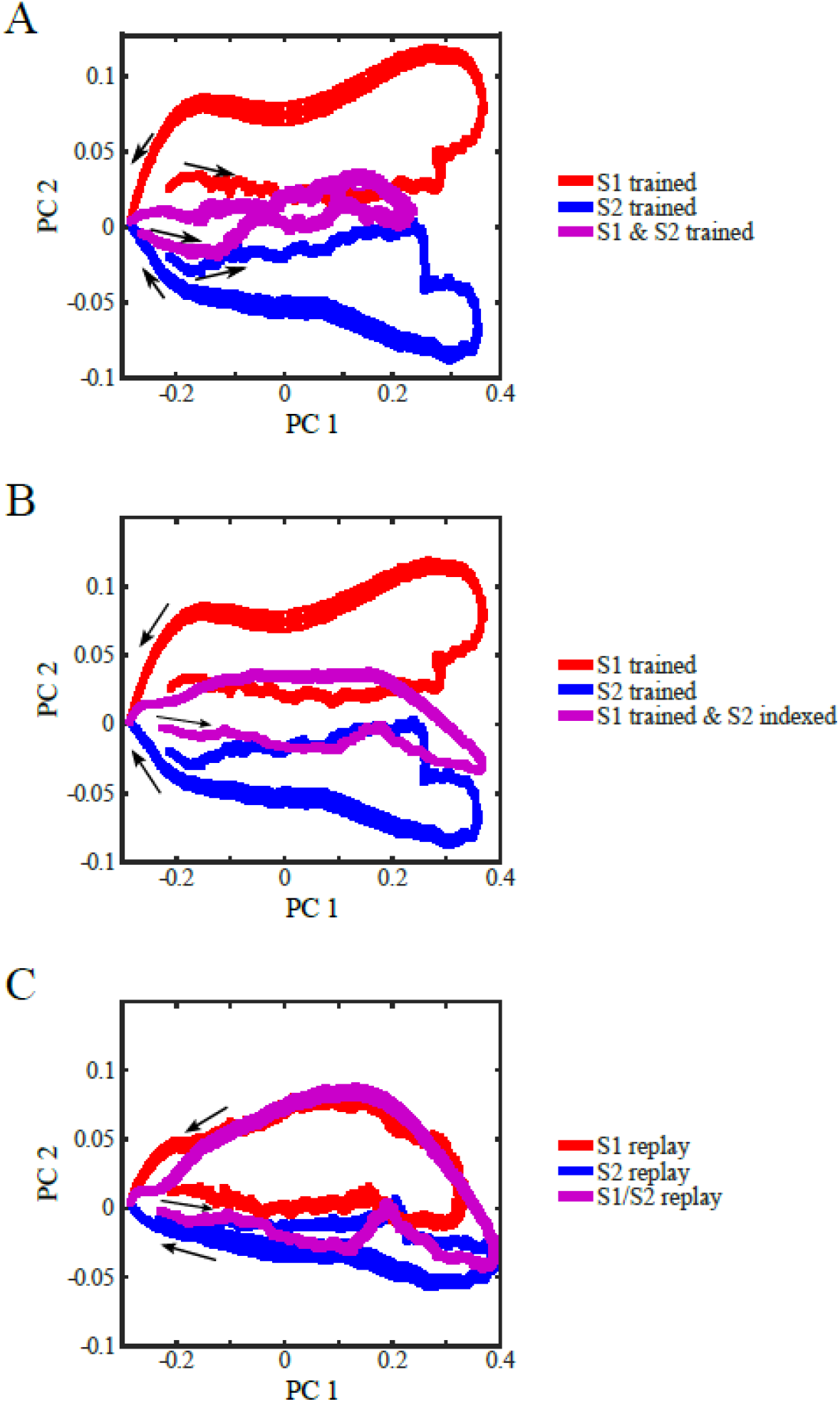
Firing-rate trajectories of memory replays during SWS are linearly separable in principal-component (PC) space. (**A**) Average firing-rate trajectories in PC space during Up states for networks trained on S1 (red), S2 (blue), or both S1 and S2 (purple) prior to sleep. (B) Average firing-rate trajectories in PC space during Up states following single-memory training (S1, red; S2, blue) and during normal hippocampal indexing of S2 in a network trained on S1 prior to sleep (purple). (C) Average firing-rate trajectories during normal indexing, grouped according to the replay probability of each Up state (>95% S1, red; >95% S2, blue; otherwise, purple).

**Supp Figure 2.**
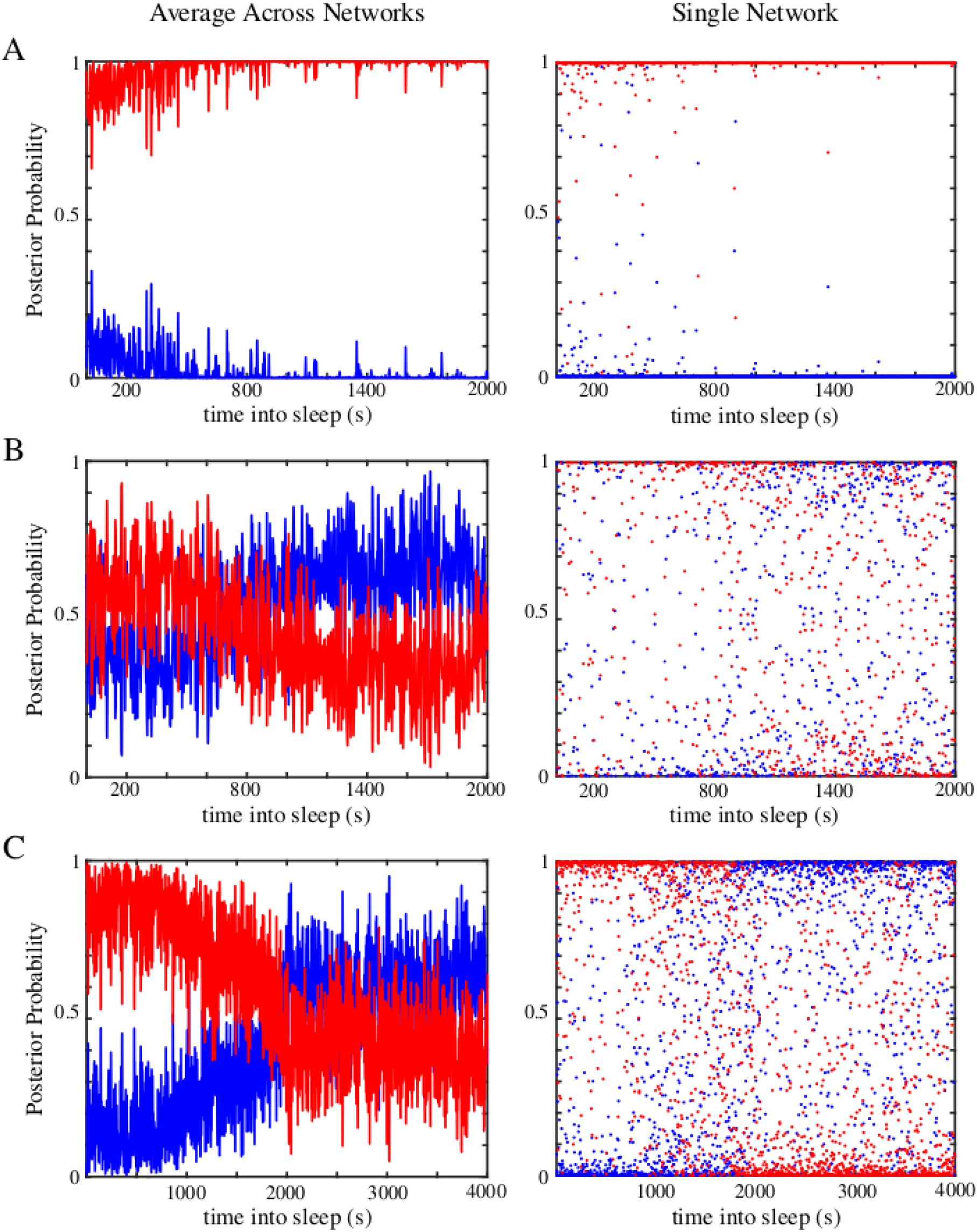
Classification of memory replay on individual UP states is robust in a single network. Averaged (left) and single-network (right) replay probabilities for (A) S1 training only, sequential training, and (C) normal hippocampal indexing. Single-network plots highlight that the majority of individual Up states are robustly classified as either S1 or S2 across all conditions.

**Supp Figure 3.**
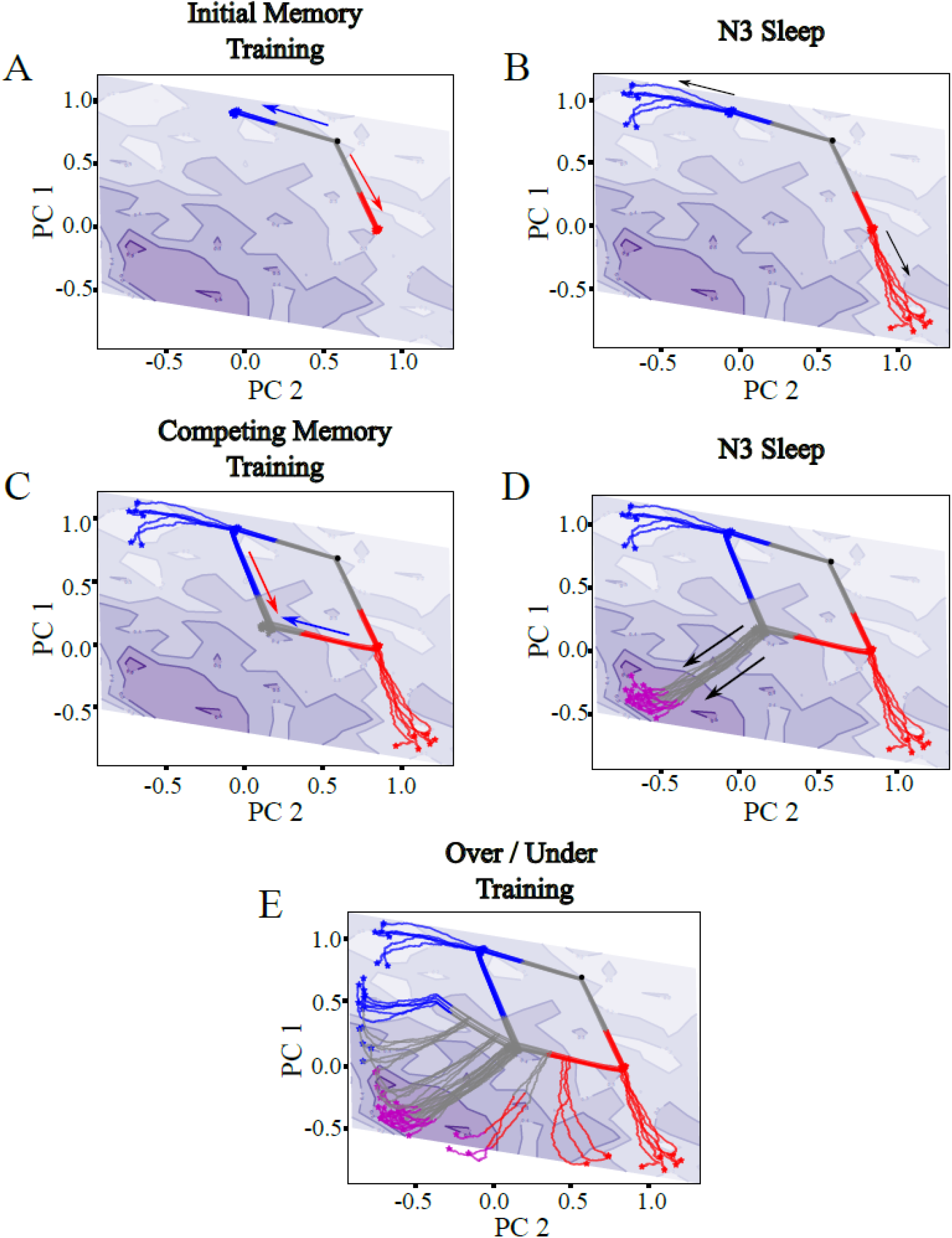
Synaptic performance landscape using a joint-memory recall metric. All panels show the evolution of the network through a dimensionality-reduced synaptic-weight space, as in Fig. 3, except that the background is now colored using a single contour plot that reflects joint-memory performance. Joint-memory performance is defined as the average of the recall performance for each memory, with the absolute difference between the two single-memory performance scores subtracted as a penalty for favoring one memory over the other. Trajectories are colored according to which memories the network can successfully pattern complete at that point in the simulation: none (gray), S1 (red), S2 (blue), or both S1 and S2 (purple).

**Supp. Fig. 4.**
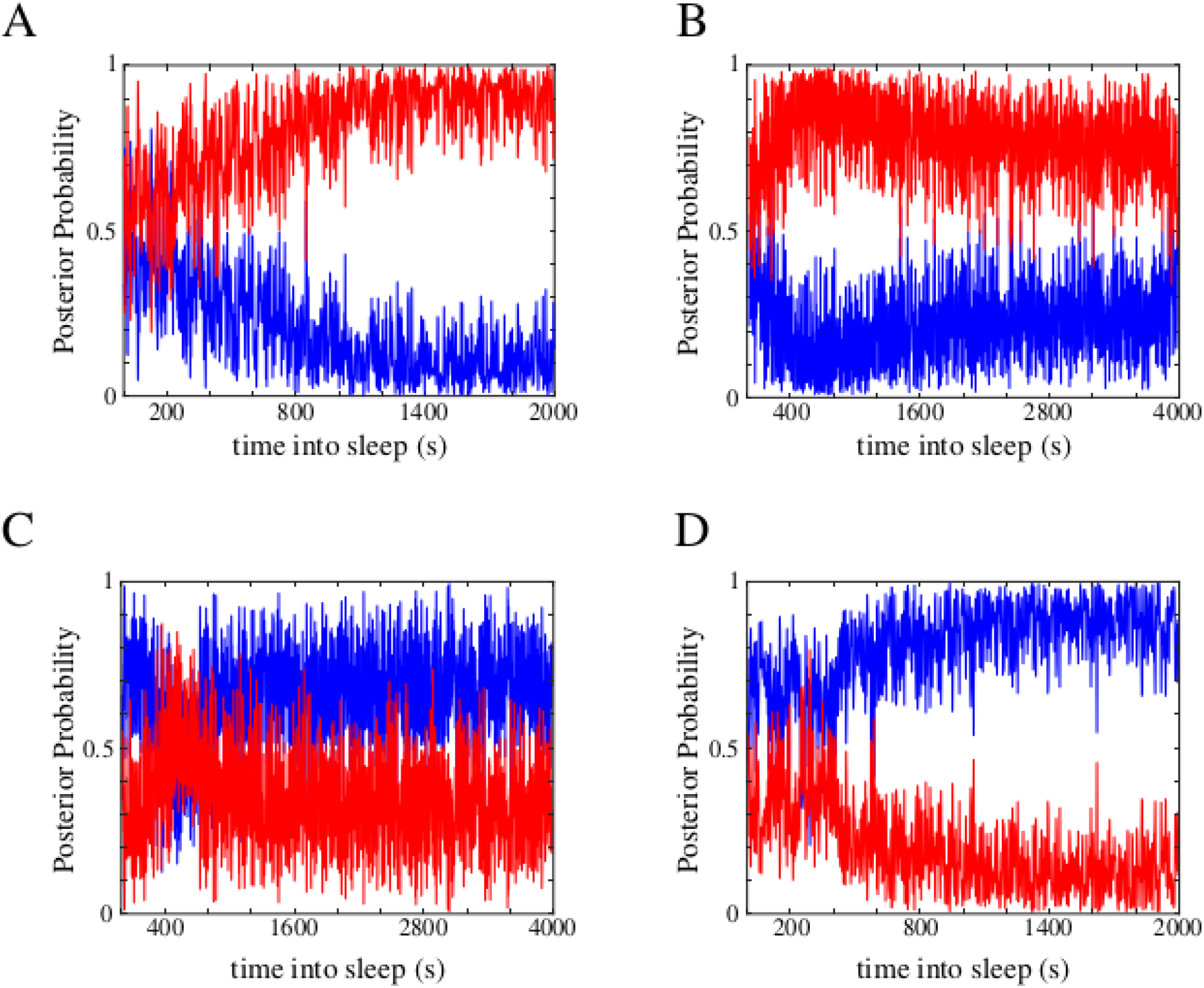
Replay probabilities during sleep following under- and over-training. (A) Following significant undertraining of S2 (Fig. 3E; Tr_1_), the network enters sleep in a mixed replay state and gradually converges to an S1-dominated replay state. (**B**) Following moderate undertraining of S2 (Fig. 3E; Tr_2_), the network diverges from an initially mixed replay state toward an S1-dominated replay state before slowly relaxing back toward a more balanced mixed state. (**C**) Following moderate overtraining of S2 (Fig. 3E; Tr_3_), the network’s replay state briefly oscillates around a mixed replay state and then remains mixed. (**D**) Following significant overtraining of S2 (Fig. 3E; Tr_4_), the network gradually converges from an initially mixed replay state toward an S2-dominated replay state.

**Supp. Fig. 5.**
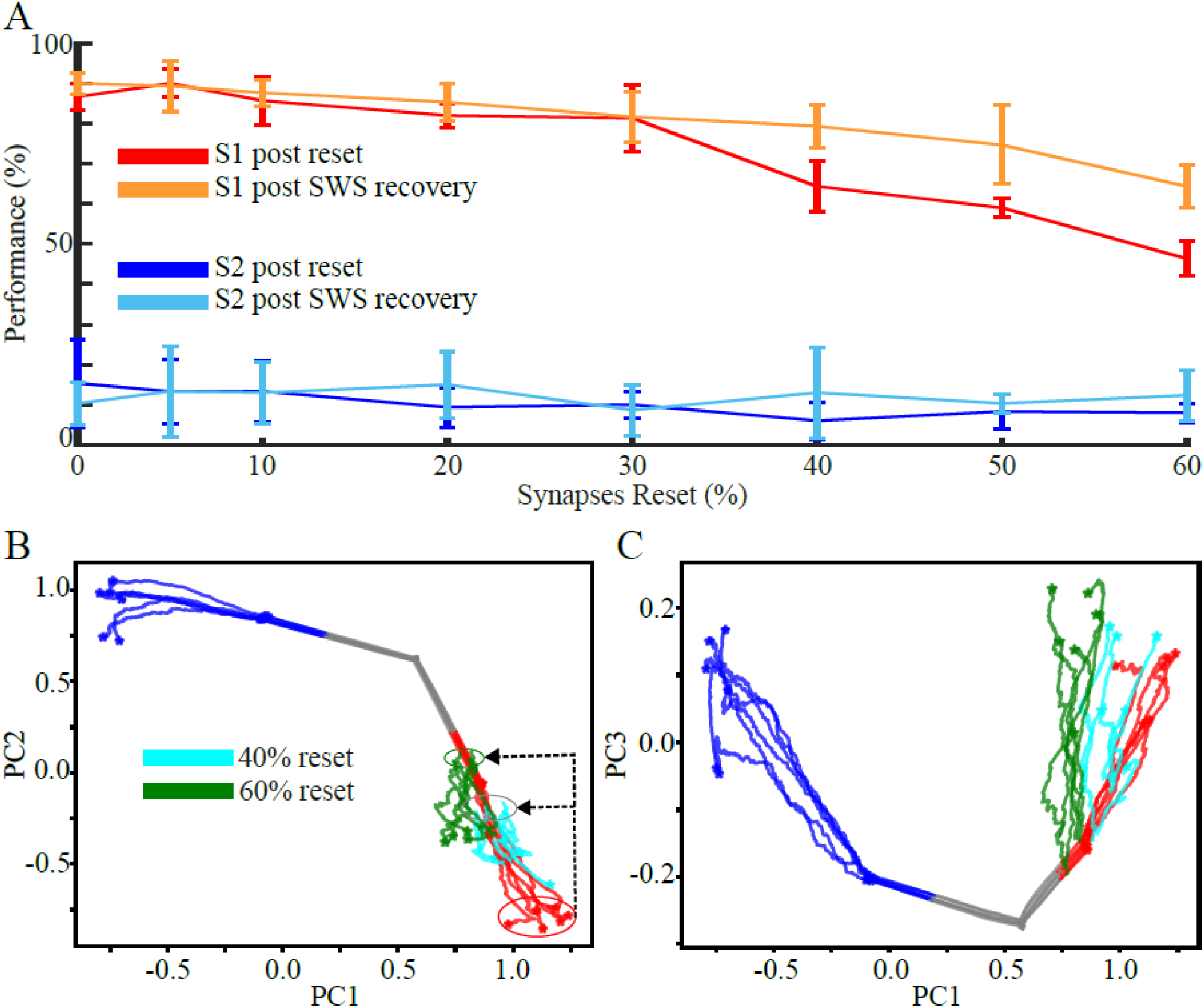
The network exhibits graceful degradation and recovery during SWS following perturbations of single-memory synaptic attractor states. (**A**) Line plots showing recall performance for S1 and S2 in a network trained on S1 during awake and consolidated with SWS, after a random fraction of synapses is reset to their initial values (S1, red; S2, blue), and following a subsequent SWS recovery period (S1, orange; S2, light blue). (**B**) Evolution of the network through synaptic-weight space after resetting 40% (cyan) or 60% (green) of synapses. Same as (B), but showing projections onto principal components 1 and 3 instead of 1 and 2.

**Supp. Fig. 6.**
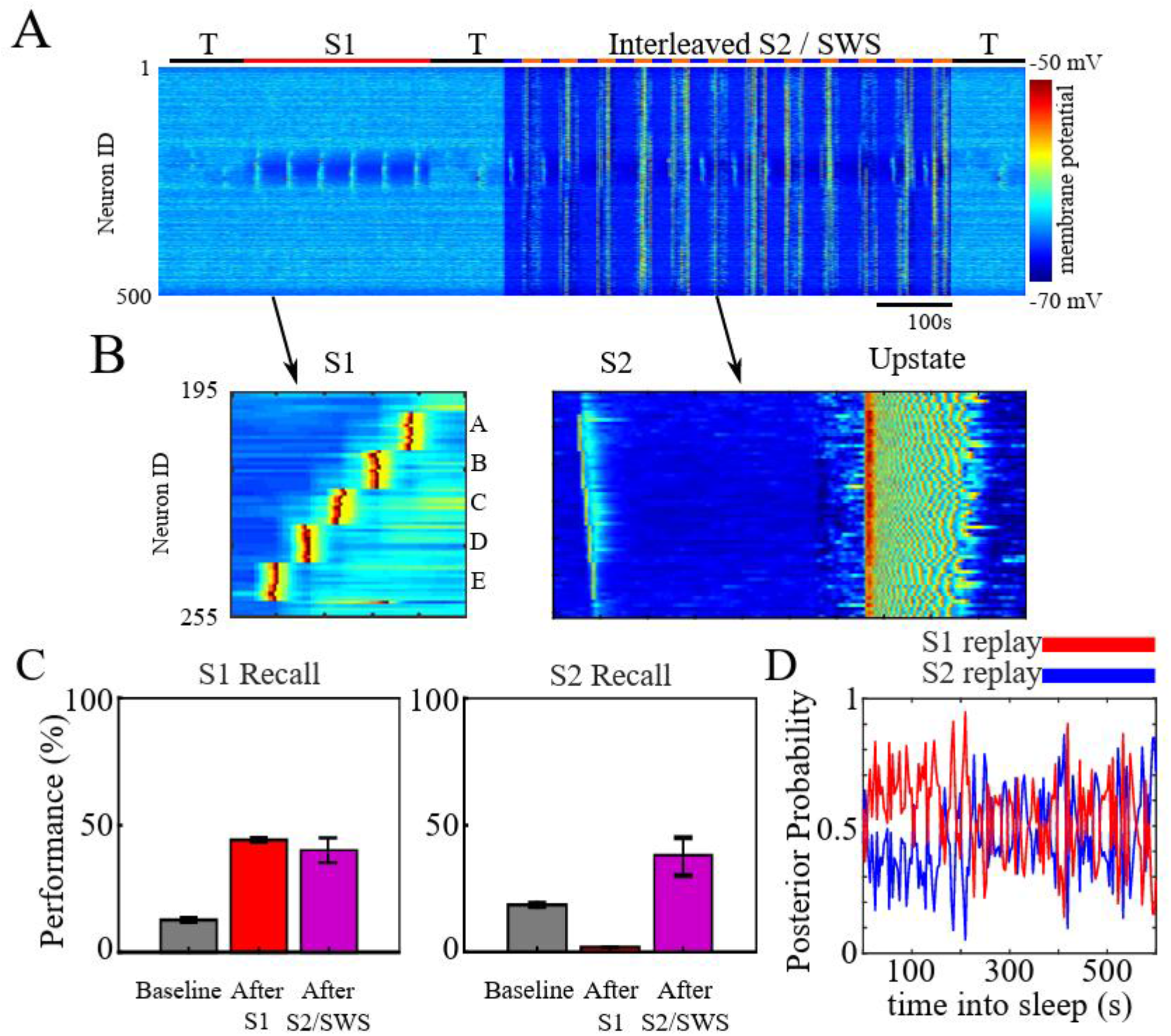
Interleaving S2 training with SWS following S1 training allows recall of both old and new memories. (**A**) Network activity during a simulation in which the network undergoes periods of testing (T; black), training on S1 (red), and 12 cycles of interleaved S2 training (blue) with SWS (orange), with each phase of the cycle lasting 25 s. (**B**) Examples of network activity during one bout of S1 training (left) and during a training-to-sleep transition, showing the final bout of an S2 training period and a single Up state at the start of the subsequent SWS period (right). (**C**) Recall performance for S1 (left) and S2 (right) shows that the network consolidates S2 without interference to S1. (**D**) Average replay probability shows that S2 (blue) replay probability initially remains below that of S1 (red) and increases to comparable levels after approximately 250 s of interleaved S2/SWS.

